# A ΦKMV ligase-dependent DNA repair mechanism that mitigates DNA-targeting nucleases

**DOI:** 10.64898/2026.09.21.753257

**Authors:** Alexis Villani, Shweta Karambelkar, Nandini Pathak, Rhea Kamath, Sutharsan Govindarajan, Bálint Csörgő, Nikolai R. Varnavski, Joseph Bondy-Denomy

## Abstract

Bacteria employ diverse DNA-targeting systems, including restriction-modification (R-M) and CRISPR-Cas, to cleave invading bacteriophage genomes. In response, phages encode counter-defense strategies that block or mitigate DNA damage. Here, we screened a panel of *Pseudomonas aeruginosa* phages against native and heterologous DNA-targeting systems and identified the *Phikmvvirus* phage genus as broadly resistant to multiple CRISPR-Cas and R-M systems. Following CRISPR-Cas12a exposure, most protospacer sequences remained genetically unchanged. However, at an intergenic protospacer, mutations accumulated with high frequency at the Cas12a cleavage site rather than within PAM or seed sequences, resembling repair-associated indels observed after genome editing in eukaryotic cells. Genetic screens to isolate Cas12a– and EcoRI-sensitized phage mutants revealed perturbations to the phage DNA ligase. A Cas12a-sensitive mutant phage was rescued by DNA ligase expression *in trans,* which was also sufficient to reverse CRISPR targeting of an unrelated phage. Together, our results support a model in which ΦKMV-like phages tolerate certain nucleases through ligase-dependent repair of nuclease-induced double-stranded breaks.

**Importance:** Bacterial resistance to antimicrobial medication is escalating, and yet new antibiotics are not readily available. Without novel antibiotics, phage therapy has emerged as a viable response to the antibiotic resistance. Ideally, phage will achieve broad host range through layered anti-defense strategies that ensure their replicative success. Here we describe a broad-acting mechanism that allows *Phikmvvirus* phages to evade nuclease targeting. Through a phage encoded DNA ligase, gp17, ΦKMV phage seems to repair at predicted cut sites, often with high fidelity but occasionally leaving scars reminiscent of NHEJ repair. Active phage DNA ligases also support nuclease evasion by a distinct phage, DMS3. These findings describe phage escape through faithful repair and identify a potentially interesting gene for phage therapy.

## Introduction

Bacteria have developed many strategies to keep bacteriophages (phages) and other genetic intruders at bay (1, 2). On the other hand, phages possess mechanisms to antagonize bacterial immune systems. Investigations into bacteria-phage warfare have revealed many novel counter-defense mechanisms (3). As examples, phages have anti-CRISPR, anti-restriction, and anti-CBASS proteins to counter specific host immune systems (3, 5). Broadly acting immune evasion strategies have also been uncovered including the nucleus-like compartment of ΦKZ-like viruses, DNA base modifications, and DNA repair, which can protect certain phages and mobile genetic elements (MGEs) from DNA-targeting immune systems (6–11). While repair pathways can enhance the emergence of PAM/seed escape mutants, whether phages can tolerate DNA cleavage through faithful repair remains incompletely understood.

Here, we designed a screen to identify “nuclease evasion” mechanisms in *Pseudomonas aeruginosa* obligately lytic phages. The screen identified phages with the ability to evade targeting by distinct CRISPR-Cas and restriction-modification nucleases in *P. aeruginosa*. Surprisingly, some lytic phages are resistant to interference by nucleases that are absent from *P. aeruginosa* genomes, (e.g. Type II-A CRISPR-Cas9 and Type V-A CRISPR-Cas12a), as well as to those common in *P. aeruginosa* (e.g. Type I-F CRISPR-Cas system and Type I restriction-modification). This suggests the presence of a broadly evasive mechanism rather than specific inhibition of a particular system. Here we uncover a ligase-dependent mechanism of CRISPR evasion in the *Phikmvvirus* genus, where repair after Cas12a cleavage results in indel-like mutations reminiscent of the NHEJ repair process in eukaryotic cells (12–13).

## Results

### Screen identifies phages broadly resistant to CRISPR and Restriction-Modification

In pursuit of understanding how phages subvert native and non-native nucleases we screened several *P. aeruginosa* lytic phages, against various CRISPR and R-M systems. DMS3 served as a model phage as it is sensitive to all systems used, while ΦKZ served as a phage that resists each system, likely due to its phage “nucleus” structure (10). Four CRISPR-Cas systems (Type I-C, I-F, II-A, and V-A) were expressed individually in strain PAO1, with two or more cognate crRNAs designed to match each phage (Fig 1A). Additionally, endogenous PAO1 Type I RM was assayed by first growing each phage in PAO1 strains lacking the R-M system, thus producing phages that will not be protectively methylated. A foreign Type II R-M enzyme (EcoRI_ was also expressed in PAO1. The differential nuclease sensitivities of several different phages will be discussed below.

**Figure 1.**
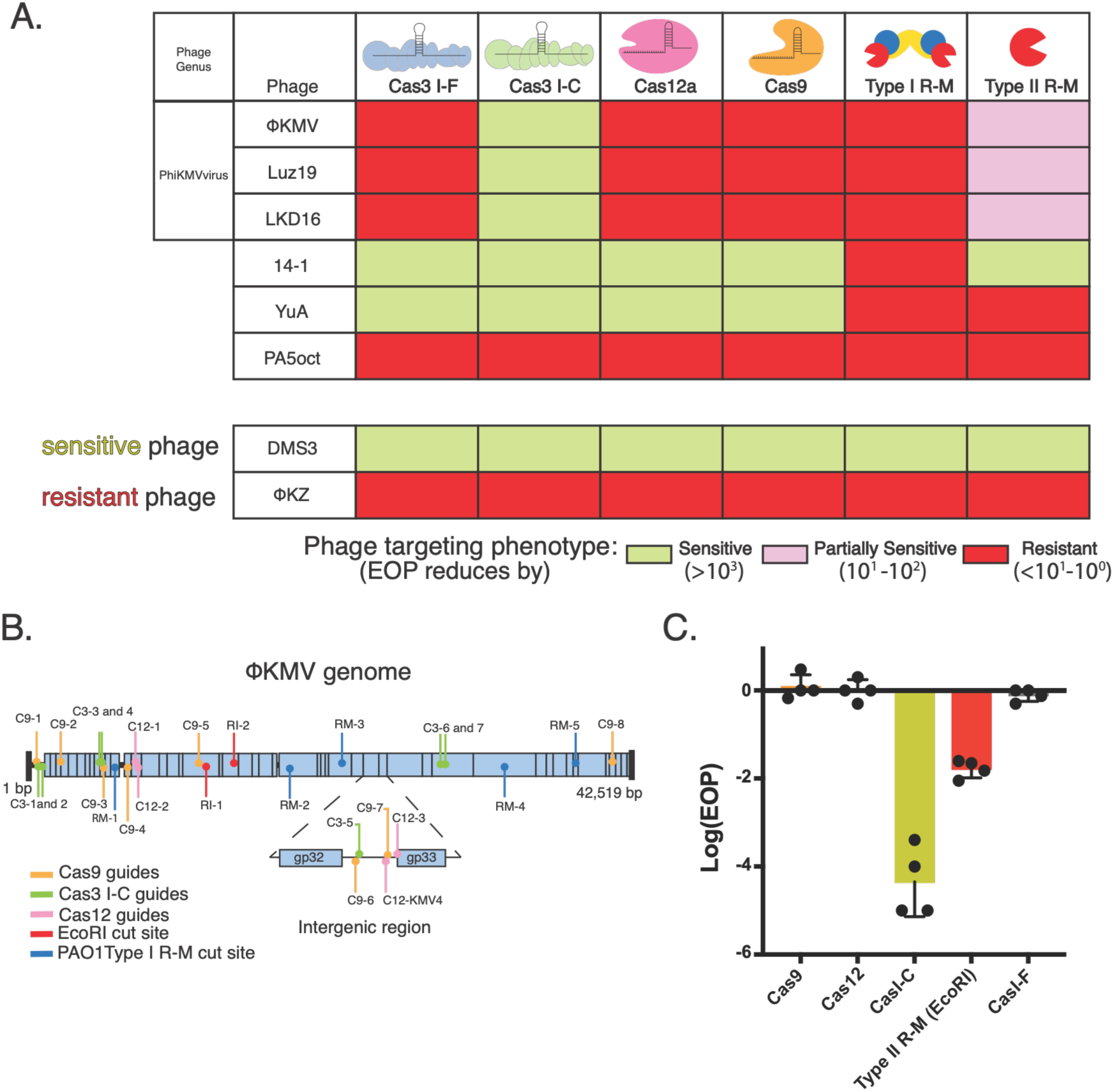
ΦKMV phages are broadly resistant to native and non-native nucleases. A. *Pseudomonas aeruginosa* phages were challenged with CRISPR-Cas and Restriction-Modification systems. The phage is denoted as sensitive to the nuclease if efficiency of plating (EOP) is reduced by >10^3^-fold. B. A schematic depicting all the protospacers used to target ΦKMV with the different DNA targeting systems. C. Log efficiency of plating (EOP) for ΦKMV, as the ratio of phage titer on a targeting strain relative to a non-targeting strain.

The *Pbunavirus* 14-1 was sensitive to most nucleases, apart from Type I R-M (Fig 1A). A recent study showed that 14-1 contains a modified genome at TG bases (15), that in theory could potentially prevent effective targeting by CRISPR-Cas systems. However, this modification does not seem to block the nucleases assayed here. Resistance to Type I R-M is notable, as the recognition motif does not contain a TG, therefore we speculate the phage bypasses targeting through another means, perhaps encoding an anti-R-M gene.

The *Yuavirus* YuA, another base-modified phage (15–17), was also sensitive to all CRISPR-Cas systems but resistant to both types of R-M systems tested (Fig 1A). YuA contains both modified thymidine and gp45, which is highly similar to anti-restriction protein ArdB (16,18). Together with the *Pbunavirus* data, this shows that modified DNA is not necessarily CRISPR-Cas resistant.

The *Wroclawvirus* Pa5oct, a jumbo phage unrelated to the ΦKZ-like nucleus-forming phages, was highly resistant to all nucleases *in vivo* (Fig S1). Incubation with EcoRI *in vitro* also failed against this phage, suggesting that the phage DNA itself is likely protected by a modification (Fig S1B). Indeed, Pa5oct was recently described to contain a hyper-modified G base and encode a functional inhibitor of END nucleases, a system that targets modified phages (12).

*Phikmvvirus* phages were fully resistant to 4 out of the 6 systems tested, with partial resistance to Type II R-M, and full sensitivity to type I-C CRISPR-Cas (Fig 1A). The location of protospacers across the phage genome is shown in Figure 1B. In addition to resisting the native immune systems of *P. aeruginosa* (type I-F CRISPR-Cas Type I-F and Type I R-M), the phages were also resistant to Cas12a and Cas9 nucleases that are not naturally found in *P. aeruginosa.* This indicated the presence of protective mechanisms effective against evolutionarily diverse host nucleases. Unlike the other resistant phages, whose mechanisms could be attributed to known factors, like base modifications or nucleus-like structures, *Phikmvvirus* resistance remained unexplained. The mechanism of nuclease avoidance by this family will therefore be the focus of this report.

### Mutation-independent escape of ΦKMV from CRISPR-Cas targeting

ΦKMV plaque formation in spot titration assays was not limited by CRISPR-Cas12a targeting (Fig 1C), which was the basic assay used in our screen. To explore whether this nuclease had any ability to target the phage or slow phage development, we conducted a more sensitive single step infection (Fig 2A). The single step infection revealed a delay and a decrease in phage release during Cas12a targeting. This replication delay suggests that the phage is cleaved or at least bound by Cas12a. To determine if ΦKMV was somehow escaping Cas12a with very high efficiency via genetic changes, we used next generation sequencing (NGS) to sequence ∼0.5-0.7 kbp across four different protospacers (two within gene 14, one within gene 33, and one intergenic, Fig 1B). No accumulation of mutations was observed within the three protospacers residing in an open reading frame, demonstrating that phages are replicating and emerging from the infection unchanged. However, at the only intergenic locus, the KMV-4 protospacer, we consistently observed a single T base deletion at position 16 of the protospacer (Fig 2B).

**Figure 2.**
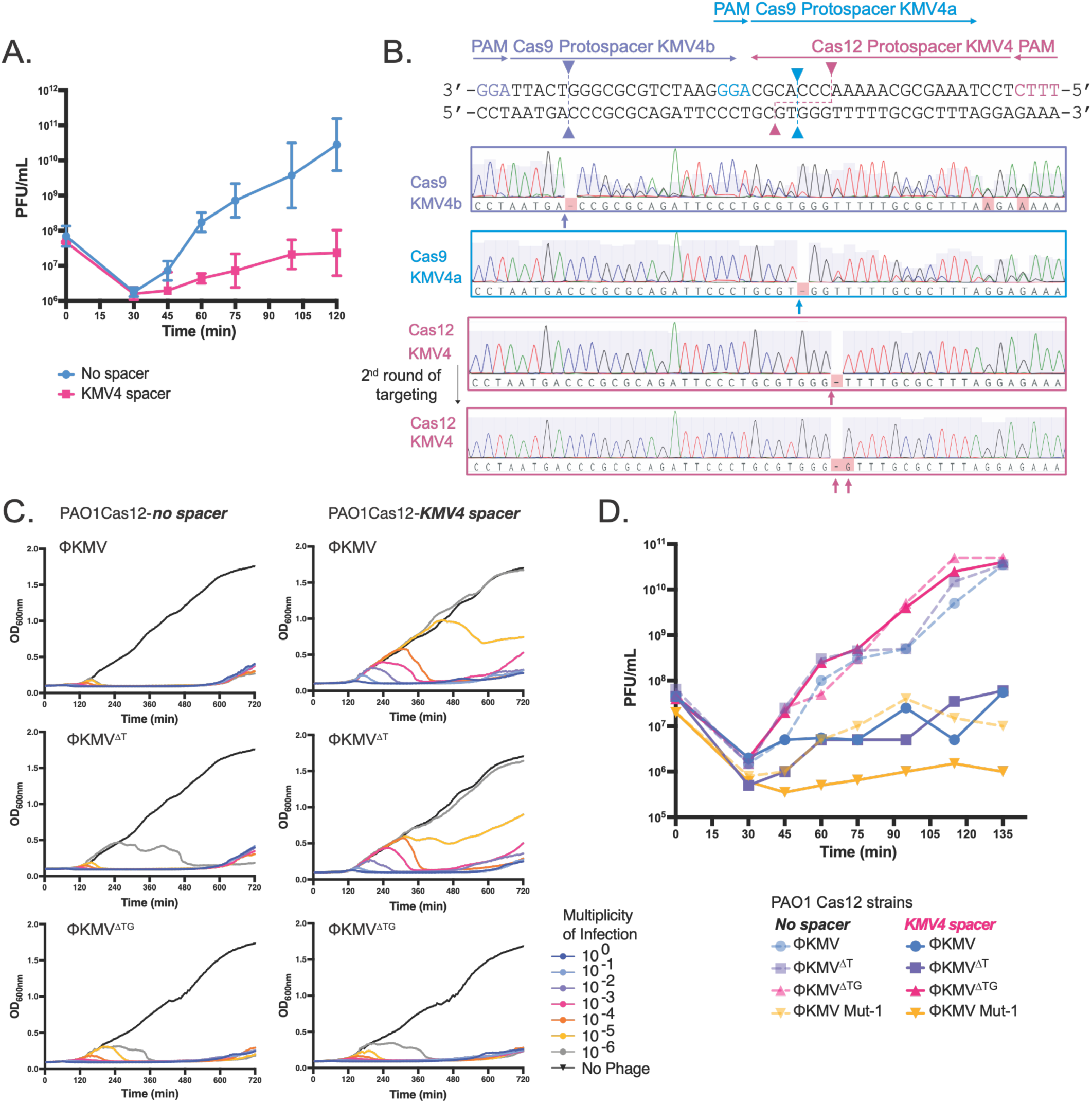
Small deletions at the cleavage site emerge in the ΦKMV upon Cas12a exposure. A. Single-step growth curves showing plaque-forming units per mL (PFU/mL) of ΦKMV over time. *Pseudomonas* PAO1 Tn7::Cas12 strains carrying either no spacer (blue) or the KMV4 spacer (pink) were infected with ΦKMV at a MOI of 0.1. Data represent three biological replicates (n = 3). B. ΦKMV intergenic sequence including the Cas9 and Cas12 protospacers. Protospacers for Cas9 KMVb (purple), Cas9 KMVa (blue), and Cas12 KMV4 (pink) are indicated by colored arrows showing spacer orientation and associated PAM sequences. Dashed arrows denote the predicted cleavage sites for each nuclease. Below, corresponding Sanger sequencing traces are shown.C. Growth curves of PAO1 Cas12 strains infected with ΦKMV, ΦKMV ^ΔT^, or ΦKMV^ΔTG^ phages across a range of multiplicities of infection (1 to 1×10^-6^). Strains expressing either no spacer or KMV4 spacer were infected with phage and optical density at 600 nm (OD_600_) was measured to monitor bacterial growth over time. Data measures technical triplicates. D. Single-step phage growth curves comparing ΦKMV and mutant derivatives, MOI = 1. Dashed lines indicate infections in a no spacer strain, while opaque lines indicate infections in a KMV4 spacer-expressing strain.

Mutations that arose in the intergenic KMV-4 protospacer were notably not in the PAM or seed position, where typical escape mutations arise, but near the Cas12a cleavage site. All individual phage plaques sequenced contain the same single T deletion at this site (Fig 2B); while NGS of parental non-targeted ΦKMV phage fails to show any point mutations in the KMV-4 protospacer region. We adapted Tracking of Indels by Decomposition, TIDE analysis, which revealed that 97.5% of phages collected from an infection with Cas12a targeting at the KMV-4 protospacer acquired a single T deletion, henceforth referred to as ΔT phage (19) (Fig S2). Targeting the related phage Luz19 with the same crRNA also yielded a mixed population of WT phages and those with a single T deletion (Fig S2). When the KMV-4 protospacer region was targeted using Cas9, which cleaves 3-5 bp upstream of the NGG PAM, mutations were also detected at the expected Cas9 cleavage site. Targeting a nearby protospacer with Cas9 also yielded single base deletions coinciding with the Cas9 cleavage site (Fig 2B). Much like Cas12a targeting, no canonical PAM escaper mutations were observed, suggesting that these are not traditional escapers but mutations that accumulate at an intergenic cleavage site.

We next attempted to determine whether the T deletion confers Cas12a escape in an adaptive manner or is a bystander result of a putative cleavage and repair event. In liquid infection experiments in the absence of CRISPR pressure, wildtype ΦKMV lysed the host at all multiplicities of infection (MOI) tested (Fig 2C). In contrast, in the presence of the Cas12a KMV4 guide, lower MOIs fail to lyse the bacterial population, further indicating that Cas12a provides modest protection to the host. When a similar liquid infection experiment was carried out with the ΦKMV^ΔT^ phage, a strikingly similar liquid infection profile to WT phage was observed, indicating that the mutant phage was still at least partially sensitive to Cas12a (Fig 2C). A single step phage burst measurement also demonstrated that Cas12a delayed the burst of this mutant phage, like WT (Fig 2D). When the ΦKMV ^ΔT^ phage was used to initiate a new infection, the emerging phage now truly ablates all Cas12 targeting. This new phage mutant, ΦKMV ^ΔTG^ carries a T→G substitution mutation, again at the Cas12 cleavage site (Fig 2B) in addition to the single bp T deletion. Ultimately, the ΦKMV^ΔTG^ phage prevents further targeting by Cas12a, independent of PAM or seed mutations. These data suggest that the single T deletion is not selected for as a Cas12a escape mutation, but rather emerges during targeting, perhaps as a flexible byproduct of cleavage and repair. Importantly, although neither intragenic nor intergenic spacers affect plating efficiency (EOP = ∼1), we speculate that these deletions can persist in intergenic regions due to the absence of selective pressure against it.

The mutations at the KMV-4 locus were reminiscent of outcomes of a typical eukaryotic Cas9/Cas12 targeting experiment, where indels arise at the cleavage site because of highly active repair pathways post CRISPR-induced DNA damage. This is in stark contrast to common phage targeting outcomes, wherein a rare pre-existing PAM or protospacer mutation gets selected under CRISPR pressure and contributes to the survival of the phage population. In the case of escape mutation selection, the efficiency of plating (EOP) is often 10^-3^ – 10^-6^, reflecting the initially low frequency of the mutants. In the case presented here, the phage may be slowed down by cleavage and repair, as shown in the single step infections, but the ultimate number of plaque forming units is unimpacted over time (EOP = ∼1). We therefore suggest that the ΔT mutation, which is resultingly in nearly the entire population is being generated *de novo* after cleavage. We next sought determine the phage gene(s) responsible for executing repair.

### Isolation of CRISPR Cas12-sensitive ΦKMV mutants

We hypothesized that the repair function is encoded in the early gene(s) of the phage. The sensitivity of ΦKMV-like phages to the Type I-C CRISPR-Cas system allowed us to use it to make deletions in the early region of the phage genome (Fig 3A). Transcriptomes of the ΦKMV-like phage Luz19 show that the first 5 kb of the genome includes 13 genes that are expressed within the first 5 min of infection (25). We designed Type I-C CRISPR-Cas guides against the individual early genes, with the expectation to create localized deletions with random boundaries. Owing to the processive activity of Cas3, we obtained a collection of phages with a wide range of deletions in the early region. Excitingly, some of these phages showed 10-1000x higher sensitivity to Cas12 targeting in traditional plaque assays compared to the WT parent phage (Fig 3A, B). Since these mutations were generated with the processive nature of the Type I-C CRISPR-Cas system, ΦKMV Mutant-1 and –2 (Mut-1 and –2, both targeted at gp8) had very different deletions (∼4.4 kbp and 2.2 kbp, respectively). Mutant-3 (Mut-3) was created by targeting gp10 and produced a ∼3.6kbp deletion (Fig 3A). Interestingly, only Mut-1 became sensitized to Cas12 targeting at the KMV-4 protospacer by ∼3-4 orders of magnitude in a standard plaque assay, as compared to wildtype phage. Mut-2 and –3 became partially sensitive to Cas12 by ∼1 order of magnitude (Fig 3B).

**Figure 3.**
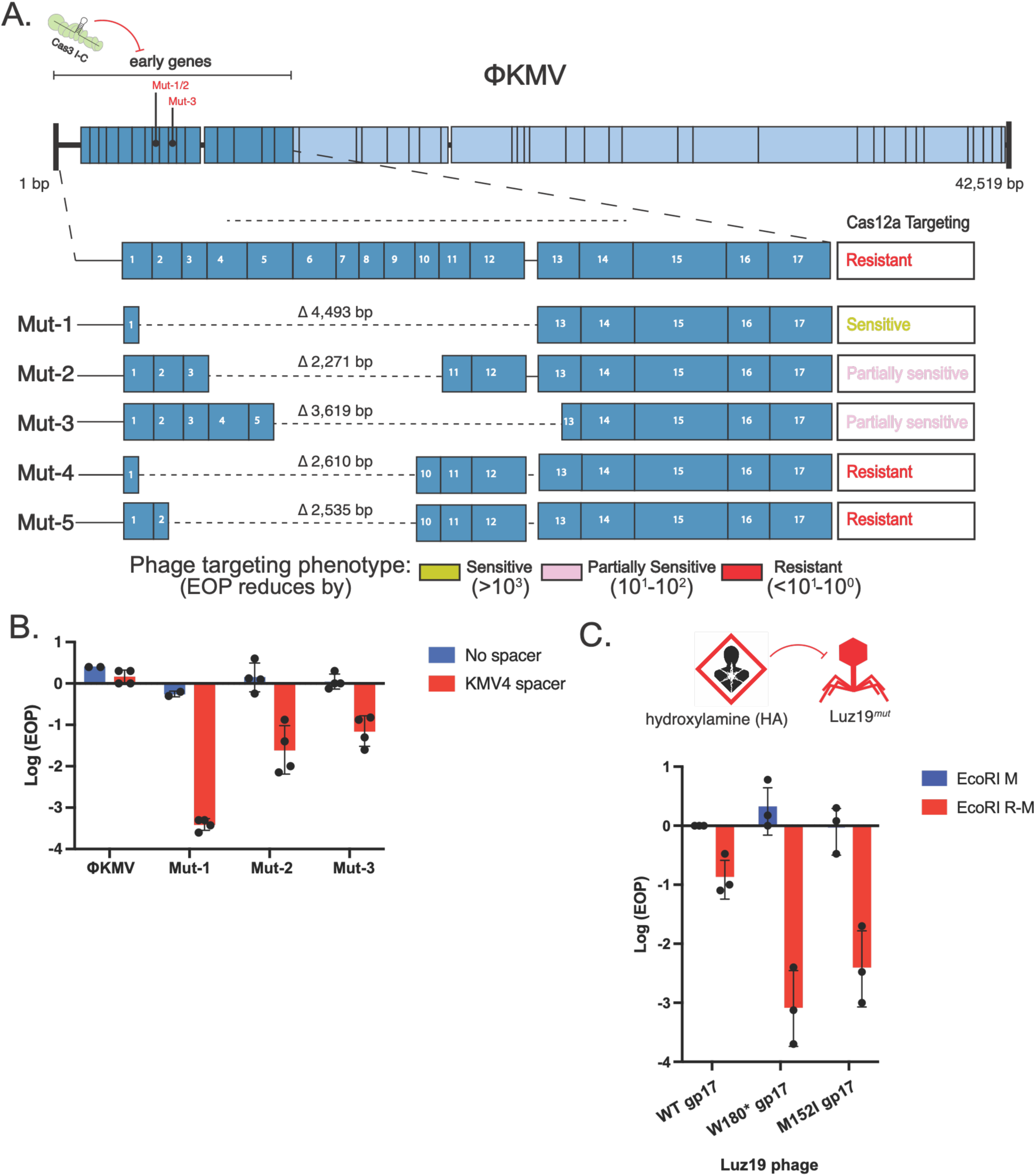
Mutant phages become sensitized to Cas12a and EcoRI. A. Schematic of the ΦKMV genome with early genes highlighted in dark blue. Individual early genes were targeted using the type I-C CRISPR-Cas system. ΦKMV mutants 1-3 were generated by targeting *gp8* or *gp10* at the indicated regions. Below, expanded views of each mutant are shown, including the corresponding deletion boundaries. Right column describes phage mutant resistance, sensitivity, or partial sensitivity towards Cas12a. B. Log-transformed efficiency of plating (EOP) for ΦKMV and mutants 1-3 on *PAO1 tn7::cas12* lawns expressing either an empty vector or a KMV4-targeting spacer. C. Schematic of the chemical mutagenesis screen (top). Luz19 phage was treated with hydroxylamine (HA), and mutants were screened for increased sensitivity to the EcoRI restriction-modification (R-M) system. The graph shows log-transformed EOP of wild-type Luz19 and a Luz19 W180* and M152I DNA ligase mutants, on PAO1 strains expressing either EcoRI methylase alone (no restriction; control) or the full EcoRI R-M system. Data represent *n* = 3 biological replicates.

Upon analysis of Mut-1-3 and other mutants, there was not a single gene whose loss could be correlated to the sensitization to Cas12a (Fig 3A), which suggested that the CRISPR sensitization was not due to deletions of any one early gene *per se.* Instead, we considered that the change in Cas12a sensitivity could emerge because of variable dysregulation of downstream genes caused by the proximal genomic deletions. Indeed, RT-qPCR assays revealed 10-1000-fold lower and delayed expression of genes immediately downstream of the deletions, which correlated well with the degree of Cas12 sensitivity (Fig S3, Fig 4A). The downstream locus encodes genes involved in phage DNA replication, including primase, helicase, DNA ligase and DNA polymerase. At the earliest timepoint of 5 minutes post infection, all three mutants showed a delay in transcript levels for gp14-17 and gp19, by 5-7 orders of magnitude. However, by 15 minutes, Mut-2 and Mut-3 catch up to wildtype expression levels, while Mut-1 continued to have a delayed expression profile by 2-4 orders of magnitude. By 35 minutes post-infection, all mutants eventually catch up to wildtype gene expression. Notably, Mut-1 shows the most significant delay in expression and is also the most sensitized to Cas12 targeting. This suggested that the downstream operon is potentially involved in DNA repair after cleavage.

**Figure 4.**
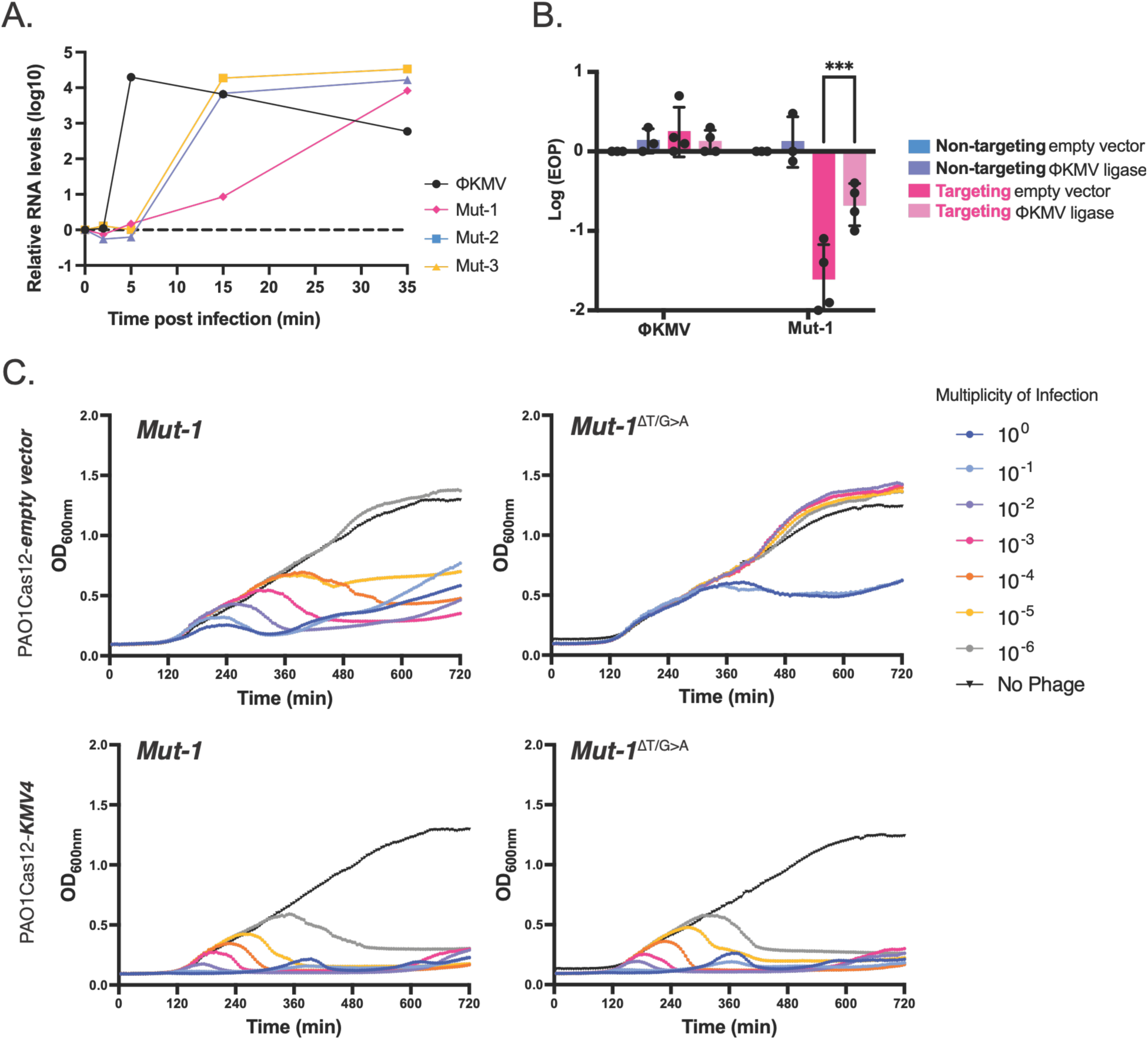
Delayed ΦKMV ligase expression in Mut-1 is correlated with increased Cas12 targeting. A. RT-qPCR measurement of gp17 (DNA ligase) over a time course infection. B. *PAO1 tn7::cas12* strains expressing either no spacer or KMV4 spacer, also expressing the ΦKMV ligase. Log EOP is shown for ΦKMV wild-type phage and Mut-1, in biological triplicates (*n=3*). (P-value = 0.0122)C. Growth curves of PAO1 Cas12 strains infected with ΦKMV Mut-1, ΦKMV Mut-1^ΔT/G>A^ (Mut-1 phage previously targeted at the KMV4 protospacer site) across a range of MOIs (1 to 1×10^-6^). Strains carried either a construct with no spacer or KMV4 spacer. Data represent technical triplicates.

### Screen for EcoRI-sensitive mutants revealed mutations in phage ligase

In parallel, we employed unbiased whole genome mutagenesis to investigate the anti-immunity factors of ΦKMV-like phages. As indicated by our screen, ΦKMV-like phages were not only resistant to most CRISPR systems tested but also to R-M systems, including partially resistant to EcoRI. A chemically mutagenized collection of ΦKMV and Luz19 was generated using hydroxylamine. Individual plaques were then picked and replica plated on targeting (EcoRI-expressing PAO1) and non-targeting host strains to identify any EcoRI-sensitized mutants. While ΦKMV generated no mutants of significance, out of a total of ∼1,300 Luz19 plaques screened, two plaques showed sensitivity to EcoRI (Fig 3C). Whole genome sequencing revealed that both phages carried mutations in the DNA ligase gene. The first mutation introduced an early stop codon at residue 180, denoted as Luz19 W180*, and the second introduced a mutation at residue 152, M152I. These data suggest a role for DNA ligase in mitigating EcoRI nuclease damage for Luz19.

During DNA replication, DNA ligase ligates Okazaki fragments (21–24). The T4 ligase has been shown to independently mediate bacterial chromosomal repair of double-stranded breaks caused by CRISPR-Cas9 (20, 28). However, the use of such DNA ligases in protecting the phage itself, has not been described to our knowledge. Based on the EcoRI mutant data that directed our attention to the phage-encoded DNA ligase, we re-examined the RT-qPCR data for ligase expression in the Cas12-sensitized mutants lacking early genes and indeed, the ligase is in the operon of genes with dysregulated expression. We observed a correlation between ligase expression and CRISPR sensitivity in the phages, with phage Mut-1, which had the most delayed ligase expression showing the maximum sensitivity to Cas12 (Fig 4A). These two independent, parallel approaches therefore coalesced on to the DNA ligase as a potentially important gene in phage resistance to EcoRI and potentially Cas12a.

### Cas12-sensitized phages showed altered repair profiles at KMV-4 locus

Targeting of wild-type ΦKMV and Luz19 phages with Cas12a at the KMV-4 locus consistently produced phages carrying a single base pair deletion within the protospacer and an EOP of 1 (Fig 2B, Fig S2). In contrast, the Cas12-sensitized phage Mut-1 exhibited an EOP of ∼10^-3^, demonstrating that most phages are unable to repair or survive this break. Surviving phages had a diversity of alleles at this locus, displaying not only a single T deletion but also 1-bp and 2-bp insertions, indicating lower fidelity repair of CRISPR-induced double-stranded breaks (Fig S4). Unlike the single T deletion, these additional mutations do enable escape from Cas12 targeting (Fig 4C).

ΦKMV is a distant relative of the well-studied phage T7 (25). Due to extensive study of the T7 phage DNA replisome, we know that the primase, helicase, and DNA polymerase encoded in this genomic region are essential components of the phage replication. However, the T7 ligase is nonessential, as host ligases can compensate (26) Similarly, Luz19 appears to tolerate ligase mutations, and we propose that this compromises its ability to efficiently repair double-stranded breaks. In Mut-1, ligase function is not impaired by mutation; rather, the delayed gene expression level prevents rapid repair of the phage during Cas12a targeting. To determine whether ligase replacement could complement Mut-1, we designed strains carrying both the Cas12a system and the spacer integrated into the bacterial chromosome. Because the crRNA was no longer expressed from a high-copy plasmid, targeting of the Mut-1 phage in a plaque assay was ∼30-fold. This system provided sufficient resolution to detect an approximately 10-fold increase in Mut-1 plaque formation upon ΦKMV ligase overexpression (Fig. 4B). Therefore, the Cas12a sensitivity in the Mut-1 phage with dysregulated DNA replication genes is largely due to lack of DNA ligase activity.

### Ligase overexpression protects DMS3 phage from CRISPR targeting

To ask whether the ligase is sufficient to rescue a phage from DNA-targeting, independently of any other ΦKMV factor, we tested it in an unrelated phage. In our screen we used DMS3 as a control phage that is broadly sensitive to DNA targeting. We designed a Cas12a guide targeting DMS3 at gp21 ΦKMV or Luz19 ligase restored DMS3 plaque forming efficiency by approximately two orders of magnitude. This rescue required ligase enzymatic activity, as neither a catalytically inactive ΦKMV ligase (K40A/R45A) nor the Luz19 W180* ligase improved phage titers (Fig 5A). NGS of the region surrounding the DMS3 protospacer revealed that overexpression of the active ligase preserved the wild-type phage sequence during targeting. These phages, therefore, remained sensitive upon retargeting by Cas12a in the absence of ligase indicating a lack of any escape mutations (Fig 5B, S5). Conversely, Cas12 targeting of DMS3 in the absence of active ligase – or upon overexpression of mutant ligase variants – reduced replicative fitness of the phage and selected for phages harboring canonical PAM and seed mutations The absence of active ligase-or upon overexpression of mutant ligase variants-. Isolation of these phages and re-exposure to Cas12a confirmed adaptive escape from Cas12 due to mutation (Fig S5). Together, these data suggest that ΦKMV and Luz19 ligases are sufficient to promote repair that mitigates CRISPR targeting, thereby influencing the balance between genome preservation and mutational escape.

**Figure 5.**
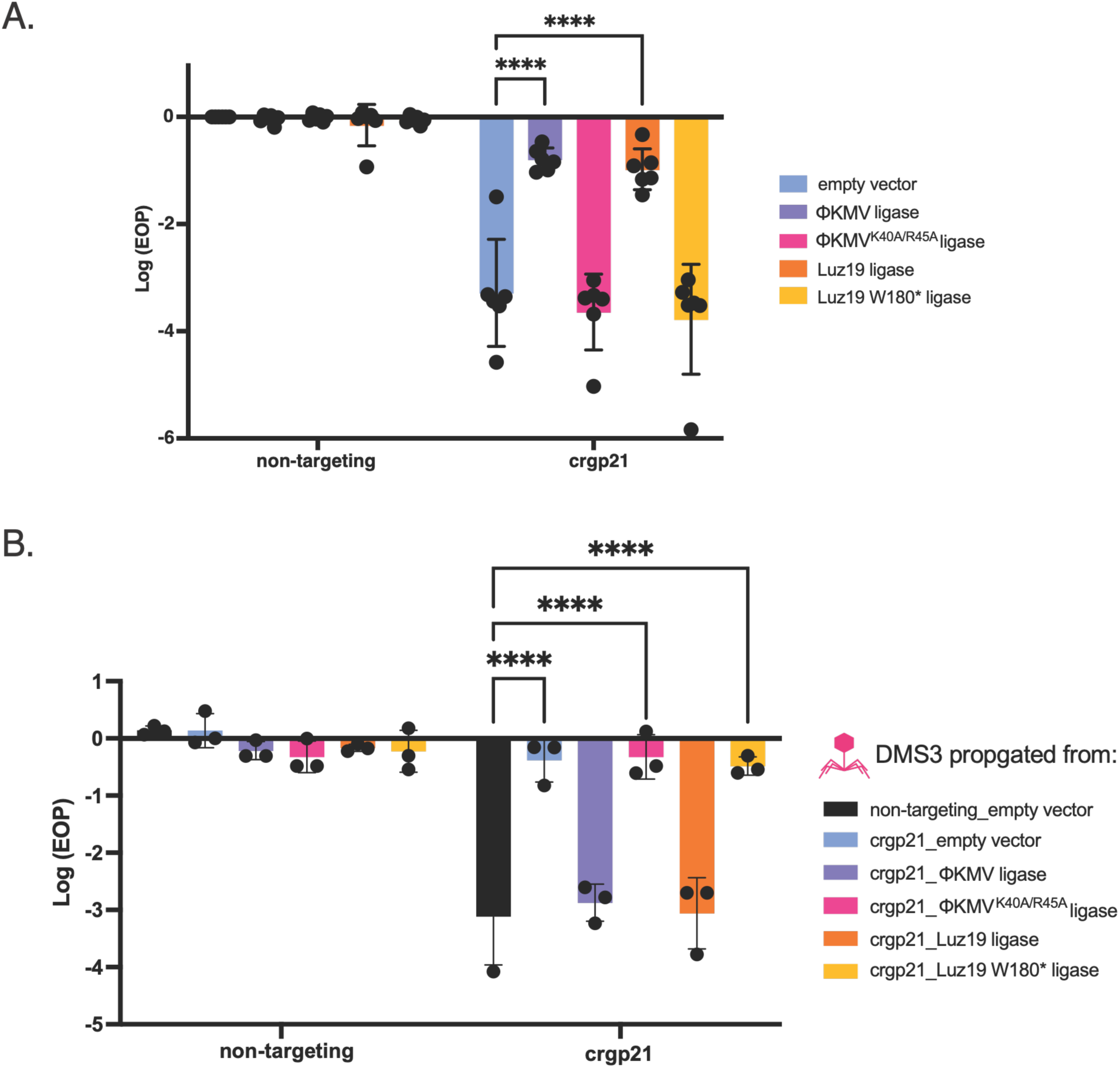
Expression of ΦKMV and Luz19 ligases rescues Cas12 targeting of DMS3. A. Full-plate infections of DMS3 phage were performed on *PAO1 tn7::cas12* strains carrying either an empty *attB* site (non-targeting) or *attB::gp21*-targeting spacer. Strains harbored plasmids expressing either no insert (empty vector), ΦKMV / Luz19 ligases, or their corresponding inactive mutants. Log efficiency of plating (EOP) is shown. Data represent *n* = 5 biological replicates. Statistical significance was determined by two-way ANOVA with multiple comparisons relative to the empty vector control: empty vector vs. ΦKMV ligase, *p* < 0.0001; empty vector vs. Luz19 ligase, *p* = 0.0007. B. DMS3 escaper phages isolated from non-targeting and targeting conditions in the presence or absence of active or inactive ligases. Individual escapers were rechallenged on Cas12-targeting strains. Data represent biological replicates (*n=2*).

## Discussion

*Phikmvvirus* phages display broad evasion of native and non-native DNA-targeting nucleases. Focusing on ΦKMV and Luz19, we found a delayed burst phenotype that is invisible to standard overnight plaque assays. Sequencing of Cas12a-targeted regions showed largely unaltered gene sequences, consistent with faithful repair or avoidance of targeting. At one intergenic locus, however, targeting reproducibly produced a single T deletion in a non-coding region, which we used as a model locus tolerant of mutation. Cas9 produced a similar outcome at this site, reminiscent of NHEJ in eukaryotic cells. This cleavage-site-localized indel did not confer escape from further Cas12 targeting, likely emerging as a byproduct of cleavage and repair. Interestingly, a phage mutant with delayed DNA ligase expression displayed very inefficient repair outcomes, which was rescued by DNA ligase expression *in trans*. Mutations in the DNA ligase of phage Luz19 also emerged, which enhanced sensitivity to EcoRI. Finally, the scar-free rescue of phage DMS3 by ΦKMV ligase expression further supports a model of cleavage followed by repair.

Some packaged phage genomes carry genomic nicks (single-strand breaks) at conserved positions, first described in phage T5 and since documented in ΦKMV-like viruses (27, 30).The function of these nicks remains unclear, but one model proposes that they facilitate packaging and are rapidly resolved during the next round of infection, potentially by the phage DNA ligase (27). Early deployment of ligase activity for this purpose could also poise the phage to repair CRISPR-Cas-induced double-strand breaks before succumbing to targeting. Canonical ligases already resolve nicks left by DNA replication, and we propose an expanded role here in nuclease mitigation via double-strand break repair: *in vitro*, ΦKMV ligase acts on both blunt and staggered breaks but is more efficient at the latter (29), suggesting it can repair breaks of distinct origins. Unlike classical CRISPR escapers, which survive through selection of pre-existing PAM or seed mutations, ΦKMV-like phages appear able to transiently tolerate nuclease cleavage through rapid, and often faithful, repair that prevents the accumulation of deleterious mutations.

## Methods

### Bacterial strains, plasmids, phages, media

All bacterial strains, plasmids, phages, and targets used in this study are described in Supplemental Tables S1 and S2. Cultures were grown on lysogeny broth (LB) agar or liquid LB at 37 °C or 30°C as indicated. In *P. aeruginosa*, PAO1, any extrachromosomal plasmids were maintained using antibiotics at the indicated concentrations: pHERD30T at 50 µg/mL. For *E.coli* strains, plasmids were maintained using pHERD30T at 15 µg/mL. Inducers were added to the media and LB agar plates at 0.1% arabinose or 0.05% rhamnose for pHERD30T plasmid.

### Construction of plasmids and strains

All transformations into *E. coli* strains, DH5alpha or XL1 Blue competent cells were prepared using the NEB (#C2987H) High Efficiency Protocol as described by the manufacturer. The following day, colonies were picked off selective plates, grown overnight at 37 °C and then mini prepped for plasmid using the Zymo^TM^ plasmid mini prep kit (Cat. No D4037). *Pseudomonas* was prepared for electroporation by washing the cells twice with 10 % glycerol in a 1:1 ratio. One milliliter of competent *Pseudomonas* cells was spun down and resuspended to 100 µL, ∼80-100 ng of plasmid DNA was incubated with cells for 30 minutes at room temperature. Cells were then electroporated on a BioRad Gene PluserXL^TM^ on a preset setting for *Pseudomonas,* then allowed to recover in a shaking incubator at 37 °C. About 50 µL of recovered cells were plated on selection plate.

### One-step growth assay

Cells were grown overnight at 37 °C in selective media. A 1:100 subculture was grown in selective and inducing media (0.3% arabinose, 0.05% rhamnose, gentamicin 50 µg/mL, 10 mM MgSO4) to an optical density (OD) of 0.3. Cells were centrifuged at 9,000 x g for 3 minutes; pellets were then resuspended in 200 µL of media. ΦKMV phage was added to cells at MOI of 0.1 and incubated on ice for 15 minutes. After incubation, 800 µL of media is added and mixed. From that mix, 50 µL is transferred to a fresh tube containing 450 µL of SM buffer and 10 µL of chloroform (timepoint 0). Cells are spun down and resuspended in 1000 µL of media, and another 50 µL is taken for the 30-minute timepoint. The bacteria-phage mixture is placed into test tubes and incubated in a water bath at 37 °C. Timepoints 0 and 30 minutes are vortexed and placed on ice. Timepoints are taken approximately every 15 minutes. The phage collection is spun down at max speed for 2 minutes. Serial dilutions of each timepoint are spotted onto the lawn of a permissive strain.

### Liquid Phage Infections

Strains were grown overnight at 37 °C in LB with respective antibiotics, a 1:1000 dilution of the overnight culture was prepared in LB with added 0.1 % arabinose and 10 mM MgSO4. We calculated the amount of phage to use by determining multiplicity of infection (MOI) from 1 to 1X10^-6^. Into each well of a 96-well plate 140 µL of the diluted culture was mixed with 10 µL of phage diluted in SM buffer. Each condition was prepared in technical triplicates. The plate reader was set to measure optical density at 600 nm (OD600), every 5 minutes at 37 °C overnight.

### Plaque assays

Strains were grown overnight at 37 °C in LB or LB with the appropriate antibiotic. 150 µL of the overnight culture was mixed with 3 mL of top agar, then poured onto LB agar plates containing 10 mM MgSO_4_ and the necessary antibiotic and inducing sugar(s) as needed for the strain. 10-fold serial dilutions of phage were prepared in SM buffer and 2 µL of each dilution was spotted on the plate. Plates were incubated at 30 °C overnight and imaged the next day.

### RT-qPCR

An overnight culture of PAO1 was diluted 1:100 into 10 mL of enriched medium (LB supplemented with MgSO^4^) and grown to mid-log phase OD_600_ 0.5. For infection assays, 450 µL of SM buffer was aliquoted into microcentrifuge tubes for each phage or mutant condition, with an additional tube containing 500 µL SM buffer serving as a no-phage control. Phage lysates (or SM buffer for the control) were combined with 500 µL of log-phase bacterial culture to initiate infection. Samples (200 µL) were collected at 0, 2, 5, 15, and 35 minutes post-infection and immediately mixed with 750 µL of DNA/RNA Shield buffer. A no-phage control sample was collected at 35 minutes. All samples were mixed thoroughly and stored at –80 °C until processing.

For RNA extraction, samples were thawed and transferred to bead-beating screw-cap tubes, and the volume of each sample was adjusted to 1.5 mL with LB medium. Cells were lysed by bead beating for three 30-second cycles. Total RNA was extracted using the Quick-RNA™ Miniprep Kit (Zymo Research; Cat. Nos. R1054/R1055) according to the manufacturer’s instructions, including an on-column DNase I digestion step. To further remove residual genomic DNA, after the manufacturer’s DNAse step, we added an additional DNAse treatment with the TURBO DNA-free™ Kit (Ambion; Cat. No. AM1907) by adding 10 µL DNase buffer and 1 µL DNase I, followed by incubation at 37 °C for 30 minutes. Subsequently, 5 µL of DNase inactivation reagent was added, and samples were incubated at room temperature for 2 minutes, then centrifuged at 10,000 × g for 1.5 minutes. The collected RNA was prepared for downstream analyses using Luna^®^ Universal One-Step RT-qPCR Kit, NEB #E3005S.

### Full plate DMS3 phage infection

*Pseudomonas* cultures were grown overnight in LB at 37 °C with antibiotics and necessary inducing solution. Serial dilutions of DMS3 stock were made to 10^-7^. Bacteria (150 µL) and phage (10 µL at various dilutions) were then incubated for 15 minutes, shaking at room temperature. Infection mix was poured in 3 mL top agar onto pre-warmed LB gentamicin 50 µg/mL, 10 mM MgSO4, 0.1% arabinose plates. Plates were incubated at 30°C overnight. Plaques were manually counted. Phage titers were calculated as follows: plaque forming units per mL (PFU/mL) = (# of plaques)*10^(dilution^ ^used)^*100

### TIDE analysis

Phage lysates were collected and then amplified by PCR using guides producing ∼700 bp fragments. Fragments size from samples was confirmed by running on a 1% agarose gel. PCR products were cleaned and concentrated using the Zymo DNA clean and concentrator kit, then set for Sanger Sequencing. A wildtype ΦKMV sequence was used as reference for TIDE analysis (19) and target sequences were input into the TIDE software.

## Acknowledgements

S.K. and A.V. designed and performed the experiments, analyzed the data and wrote the manuscript. R.K., N.P., N.V. and S.G. performed phage plaque and microplate liquid assays and analyzed the data. B.C. designed and constructed the Cas3 crRNA vectors. S.S. performed NGS and analyzed the data. We would like to thank Sukrit for his help with sequencing and answering questions. J.B.D. conceived and supervised the study, designed the experiments, acquired the funding and wrote the manuscript.

J.B.-D. is supported by the National Institutes of Health (R01 AI167412) and support from the Bowes Biomedical Fellowship and UCSF. A.V. was supported by the National Science Foundation (NSF)

---

**Supplementary Table S1.**
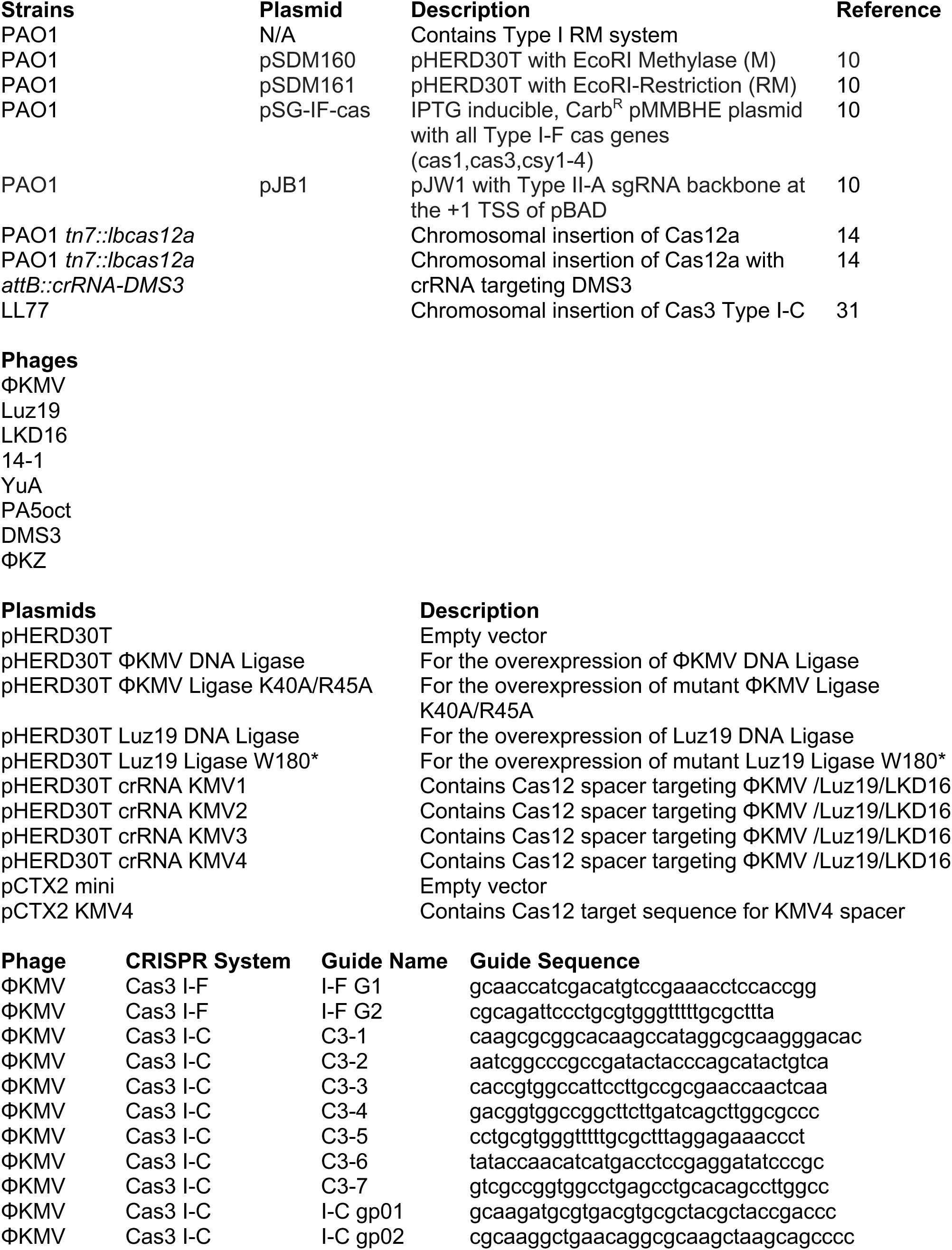
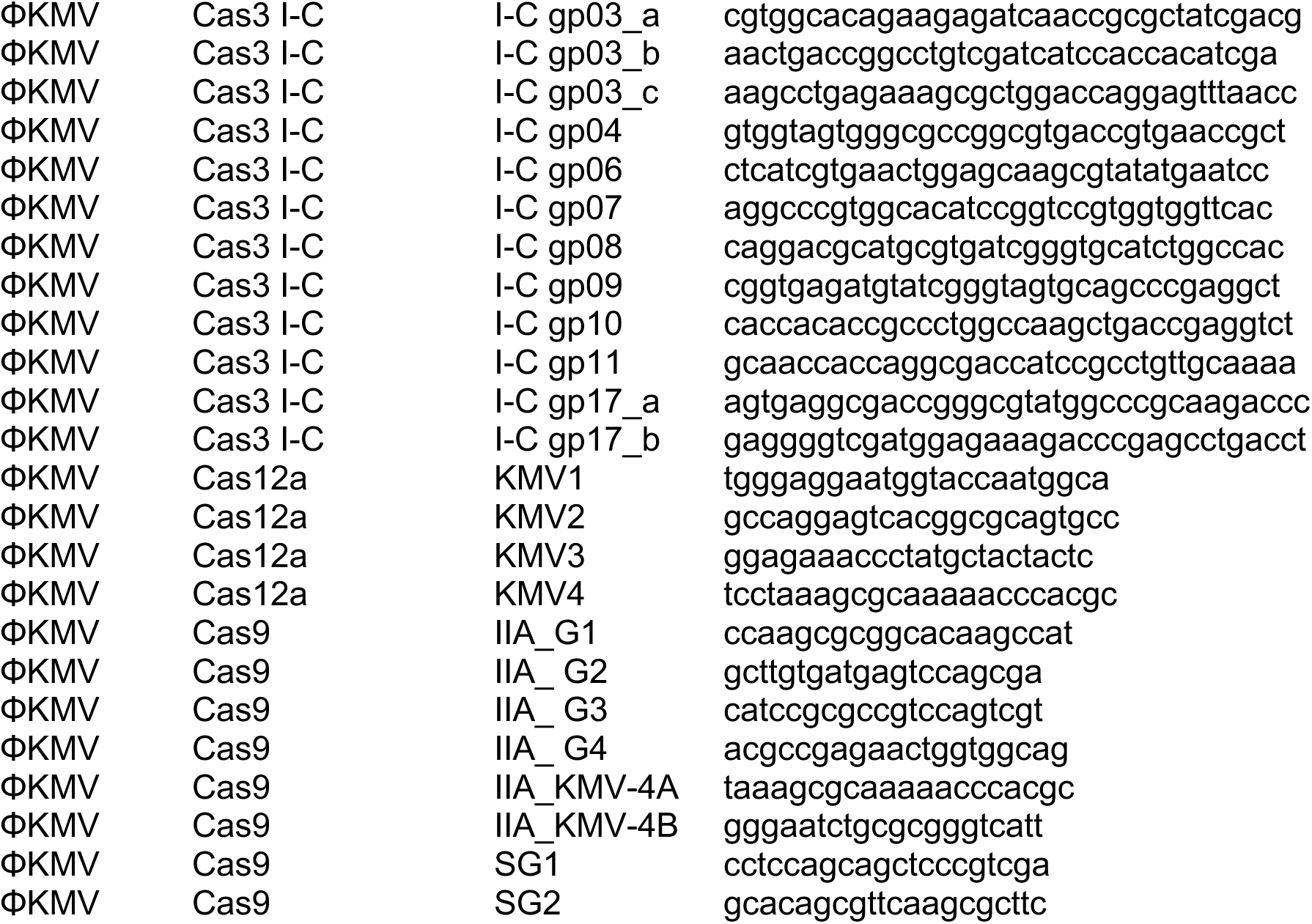
Strains, phages, plasmids, and crRNAs used in this study.

**Supplementary Table 2.**
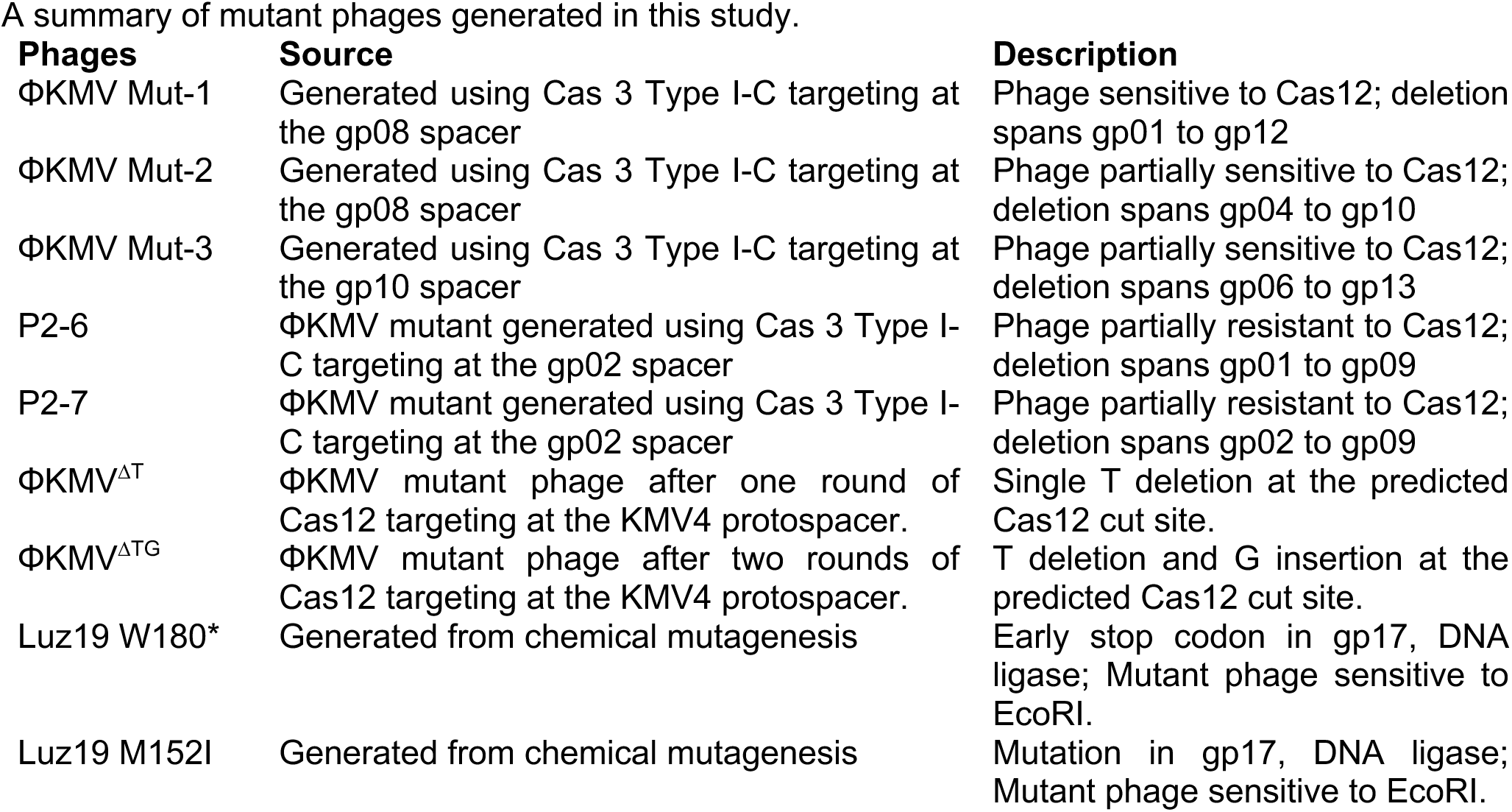
Mutant Phages.

**Supplementary Figure 1.**
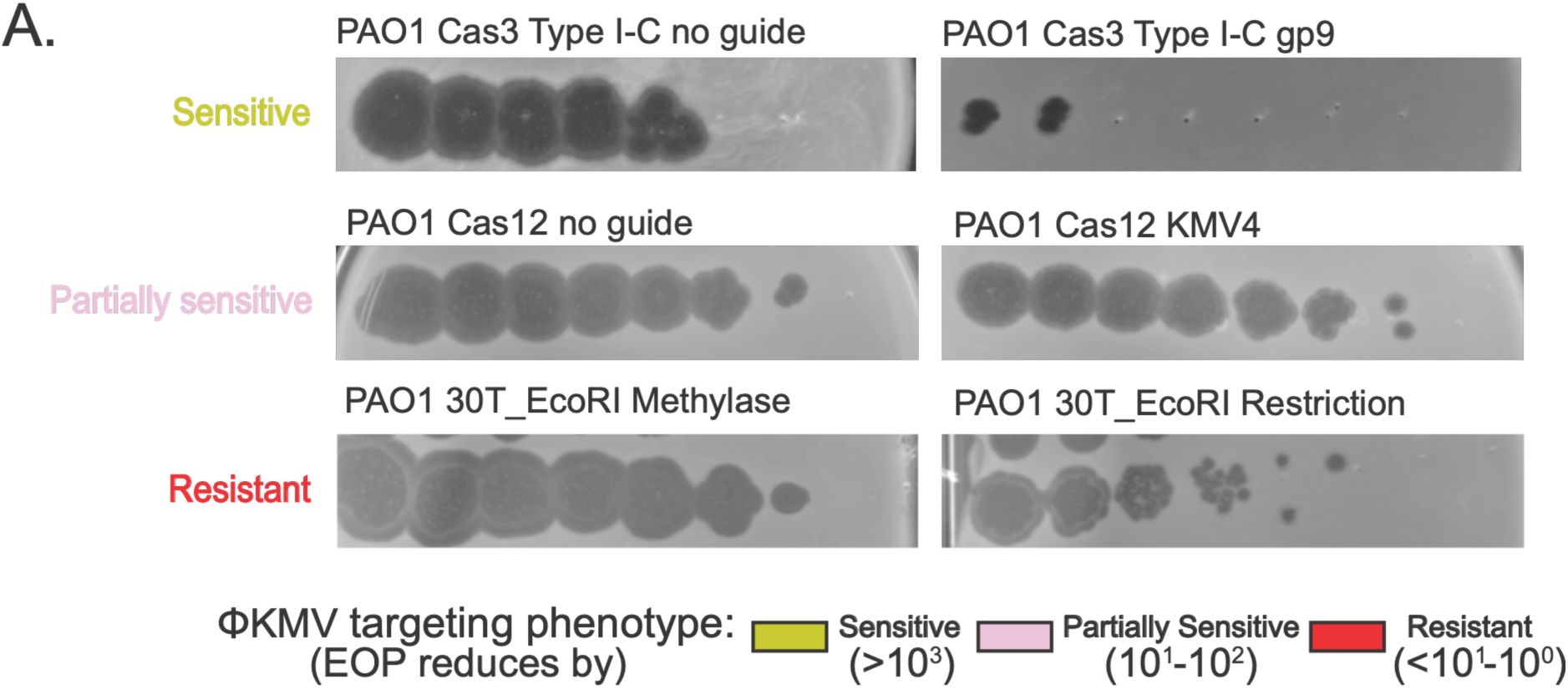
Examples of ΦKMV spot assays used to determine phage sensitivity. A. Spot Assays measure phage replication of ΦKMV spotted on the lawn of nontargeting and targeting strains. On the left side of the panel, phage phenotype is highlighted as sensitive, partially sensitive, and resistant. Similar spot assays were used to generate the summarized phage phenotypes in Figure 1A.

**Supplementary Figure 2.**
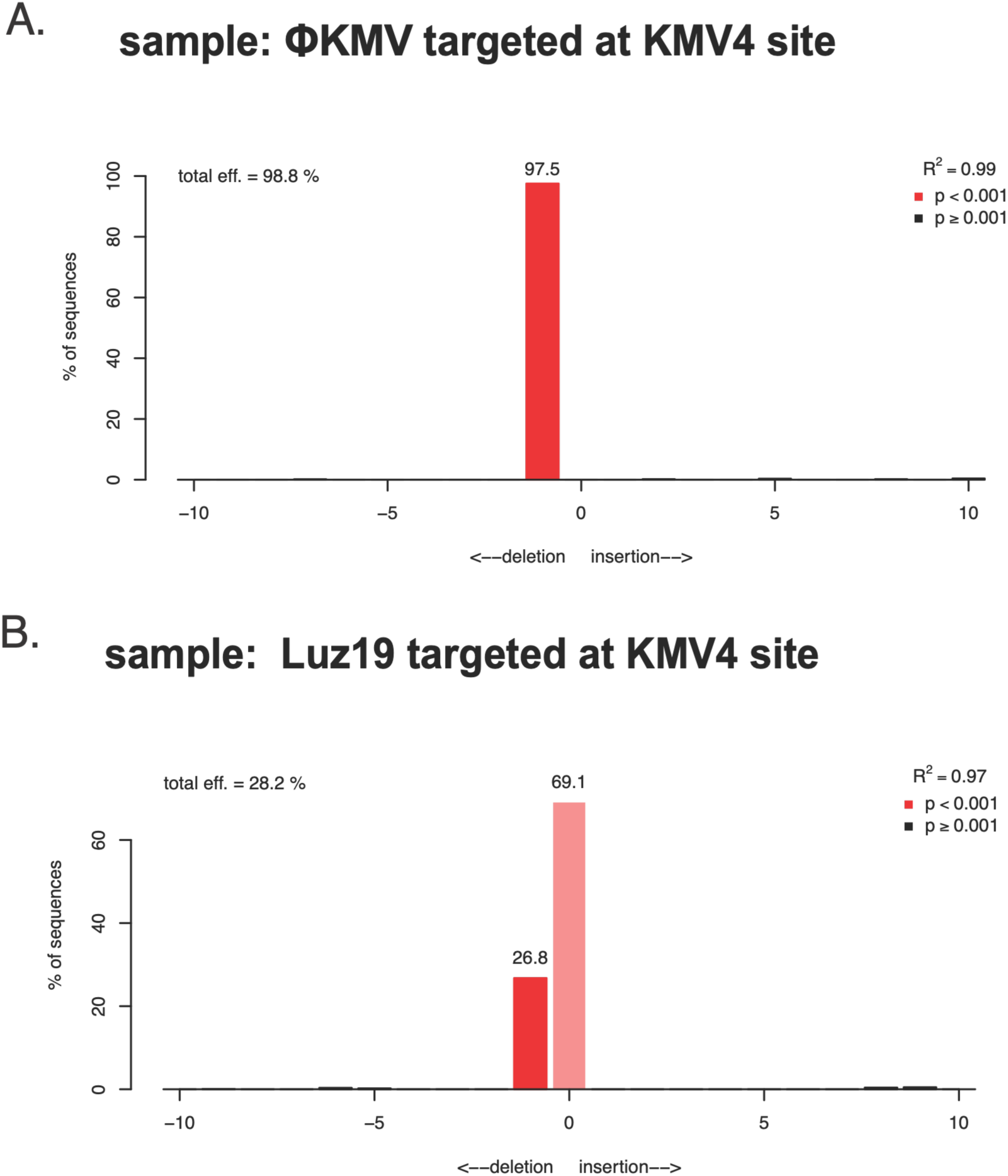
TIDE analysis of a population of ΦKMV and Luz19 phages propagated on PAO1 Cas12 strain with KMV-4 spacer. A. TIDE analysis of ΦKMV phage targeted at the KMV4 site with CRISPR Cas12. On the y-axis the graph depicts percentage of sequences, total eff or total efficiency is the percentage of sequences that contain indels. The x-axis at 0 represent sequences without indels; +/− n (number on the x axis is the number of indels present in the sequences input). The red bar represents 99.1% of sequences input contain a single deletion at a p value < 0.001. B. TIDE analysis of Luz19 phages.

**Supplementary Figure 3.**
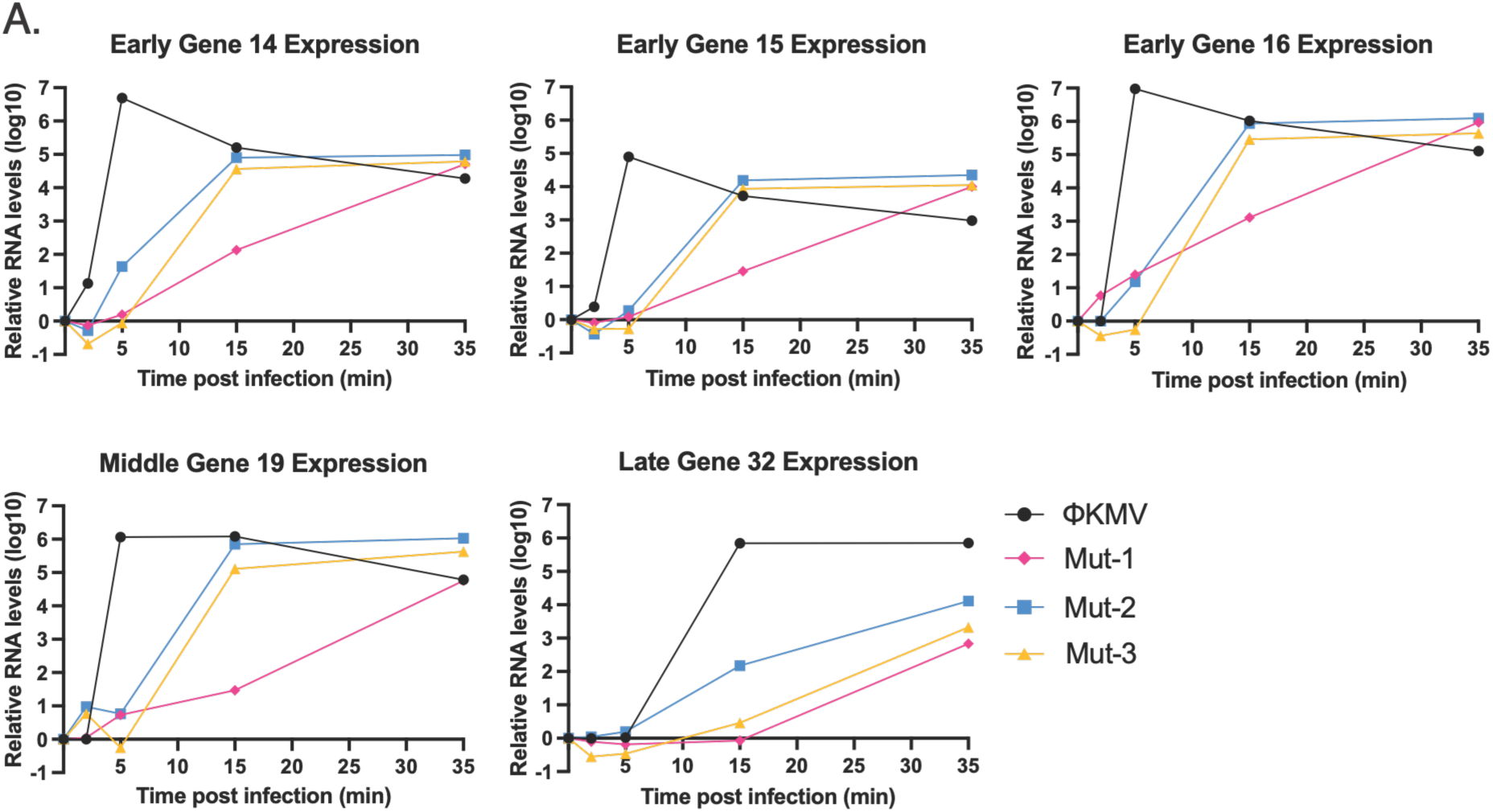
Cas I-C mutants and gene expression profiles of ΦKMV and mutants. A. Gene expression profiles of middle and late genes measured by RTPCR for ΦKMV (black), and Mut1-3 (yellow, pink, and blue, respectively). The y-axis measures the expression of the target gene against a reference gene. The X-axis measures time post infection with the indicated phages.

**Supplementary Figure 4.**
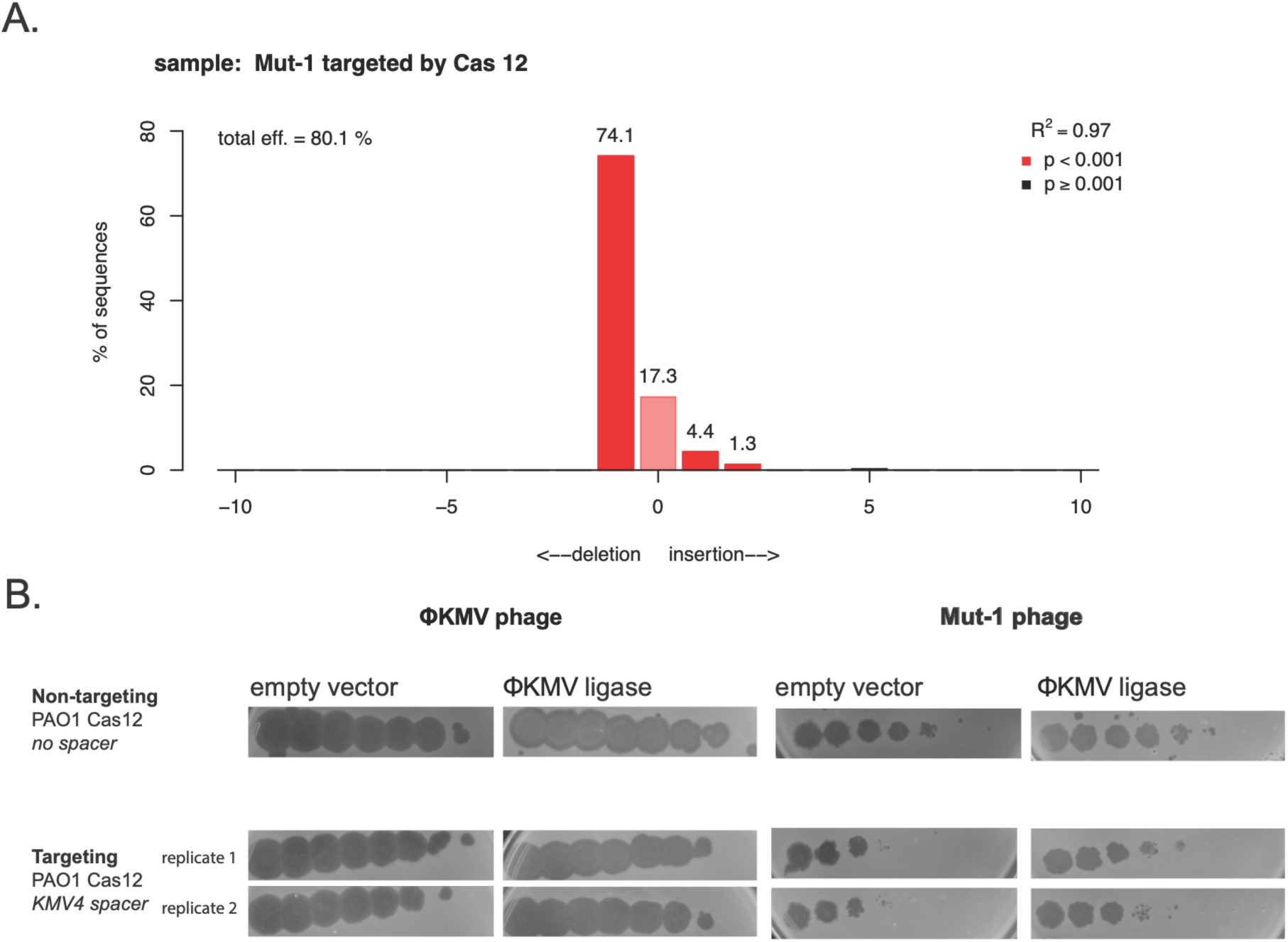
Mut-1 phage which generates many indels after targeting, is complemented by ΦKMV ligase expression. A. TIDE analysis on ΦKMV Mut-1 phage targeted at the KMV4 protospacer by Cas12. A total efficiency of 80.1% represents the population of sequences with indels. The decomposition of those indels shows 17.3% include no indels, 74. 1% include a single deletion, 4.4% included a single insertion, and 1.3 % included 2 insertions. A wildtype ΦKMV sequence was used as the control reference for TIDE analysis. B. Spot Assays measure phage replication of ΦKMV (on the left) spotted on the lawn of nontargeting and targeting strains, with the extrachromosomal expression of empty vector or active ligase. On the right side of the panel, Mut-1 phage is spotted against the same strains. This data highlights the improved replication of Mut-1 when active ΦKMV ligase is expressed during targeting. The targeting strain spot assays are biological replicates.

**Supplementary Figure 5.**
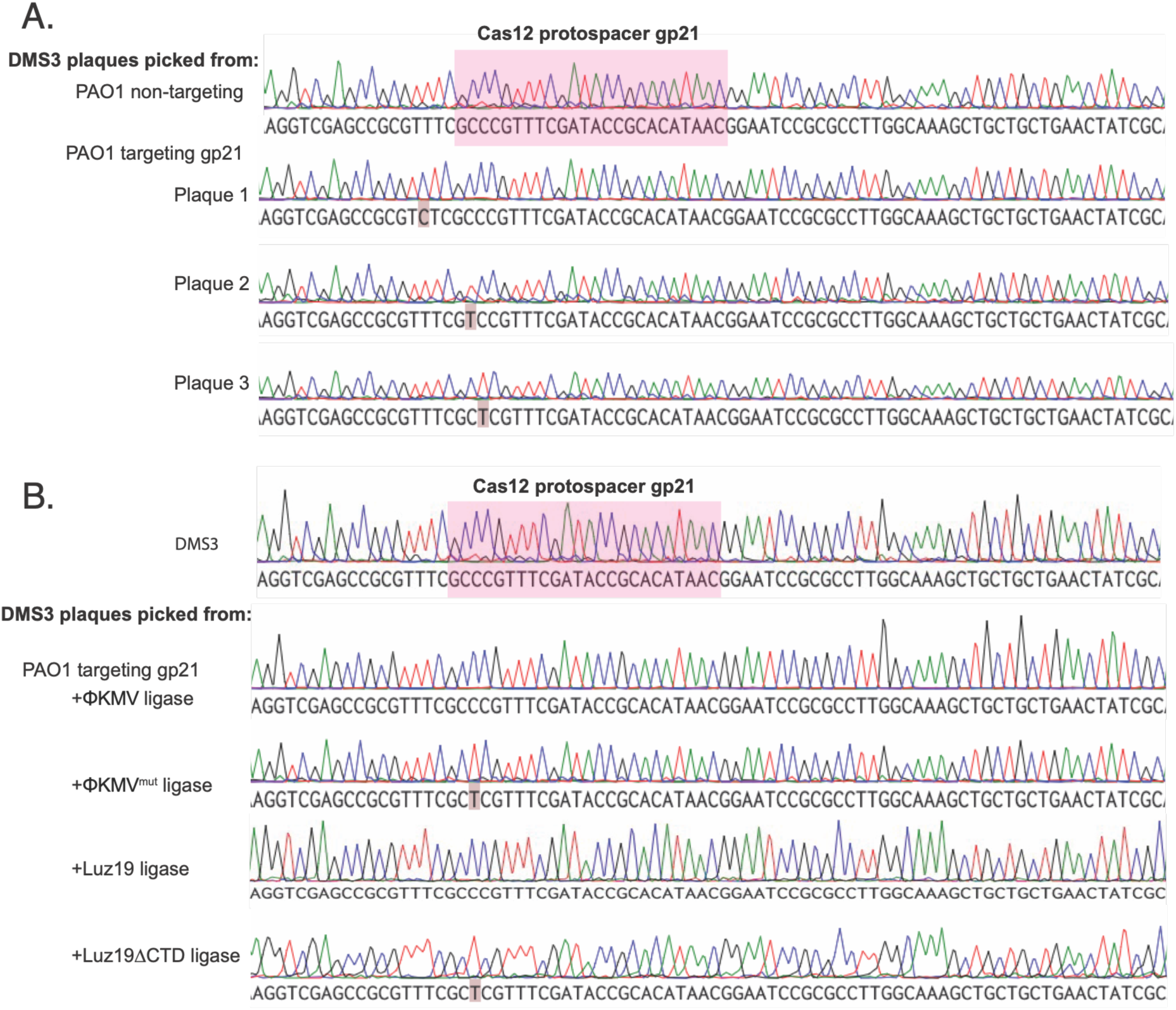
Sanger Sequence traces reveal ΦKMV ligase is sufficient to help DMS3 escape targeting without generation of indels. A. Sanger Sequencing of region surrounding the gp21 Cas12 protospacer in DMS3. The first read is of a wildtype DMS3 phage picked on a non-targeting PAO1 strain, highlighted in pink is the sequence of the protospacer. The bottom 3 reads are individual plaques picked of DMS3 propagated on a gp21 targeting strain. The sequences here show accumulation of PAM and seed mutations. The PAM sequence which is not highlighted here is the TTTN sequence downstream of the protospacer seen in pink. B. Like Fig S5A, the top sequence is of a wildtype DMS3 phage and the region around the protospacer in pink. The bottom sequences are of DMS3 single plaques picked off a lawn of PAO1 Cas12 targeting (gp21) strains with the added vector of ΦKMV ligase, ΦKMV catalytically dead ligase (K40A/R45A), Luz19 ligase, and Luz19 ΔCTD ligase. Over expression of active ligase produced perfect sequence reads, while the inactive ligases produced mutations in the seed region of the sequence.

## References

1. Wein, T., Sorek, R. Bacterial origins of human cell-autonomous innate immune mechanisms. Nat Rev Immunol 22, 629–638 (2022). 10.1038/s41577-022-00705-4

2. Bernheim, A., Sorek, R. The pan-immune system of bacteria: antiviral defence as a community resource. Nat Rev Microbiol 18, 113–119 (2020). 10.1038/s41579-019-0278-2

3. Murtazalieva, K., Mu, A., Petrovskaya, A. The growing repertoire of phage anti-defence systems. Trends Microbiol 32, 1212–1228 (2024). 10.1016/j.tim.2024.05.005

4. Niault, T., van Houte, S., Westra, E., Swarts, D.C. Evolution and ecology of anti-defence systems in phages and plasmids. Curr Biol 35, R32–R44 (2025). 10.1016/j.cub.2024.11.033

5. Bondy-Denomy, J., Pawluk, A., Maxwell, K.L., Davidson, A.R. Bacteriophage genes that inactivate the CRISPR/Cas bacterial immune system. Nature 493, 429–432 (2013). 10.1038/nature11723

6. Wu, X., Zhu, J., Tao, P., Rao, V.B. Bacteriophage T4 escapes CRISPR attack by minihomology recombination and repair. mBio 12 (2021). 10.1128/mbio.01361-21

7. Roy, D., Huguet, K.T., Grenier, F., Burrus, V. IncC conjugative plasmids and SXT/R391 elements repair double-strand breaks caused by CRISPR-Cas during conjugation. Nucleic Acids Res 48, 8815–8827 (2020). 10.1093/nar/gkaa518

8. Hossain, A.A., McGinn, J., Meeske, A.J., Modell, J.W., Marraffini, L.A. Viral recombination systems limit CRISPR-Cas targeting through the generation of escape mutations. Cell Host Microbe 29, 1482–1495 (2021). 10.1016/j.chom.2021.09.001

9. Wang, S., Sun, E., Liu, Y., Yin, B., Zhang, X., Li, M., Huang, Q., Tan, C., Qian, P., Rao, V.B., Tao, P. Landscape of new nuclease-containing antiphage systems in Escherichia coli and the counterdefense roles of bacteriophage T4 genome modifications. J Virol 97, e00599–23 (2023). 10.1128/jvi.00599-23

10. Mendoza, S.D., Nieweglowska, E.S., Govindarajan, S., Leon, L.M., Berry, J.D., Tiwari, A., Chaikeeratisak, V., Pogliano, J., Agard, D.A., Bondy-Denomy, J. A bacteriophage nucleus-like compartment shields DNA from CRISPR nucleases. Nature 577, 244–248 (2020). 10.1038/s41586-019-1786-y

11. Weigele, P., Raleigh, E.A. Biosynthesis and function of modified bases in bacteria and their viruses. Chem Rev 116, 12655–12687 (2016). 10.1021/acs.chemrev.6b00114

12. Cisneros-Aguirre, M., Ping, X., Stark, J.M. To indel or not to indel: factors influencing mutagenesis during chromosomal break end joining. DNA Repair 118, 103380 (2022). 10.1016/j.dnarep.2022.103380

13. Song, B., Yang, S., Hwang, G.H., Yu, J., Bae, S. Analysis of NHEJ-based DNA repair after CRISPR-mediated DNA cleavage. Int J Mol Sci 22, 6397 (2021). 10.3390/ijms22126397

14. Marino, N.D., Pinilla-Redondo, R., Bondy-Denomy, J. CRISPR-Cas12a targeting of ssDNA plays no detectable role in immunity. Nucleic Acids Res 50, 6414–6422 (2022). 10.1093/nar/gkac462

15. Yee, WX., Lee, YJ., Makarova, K.S. et al. END nucleases are antiphage defence systems targeting multiple phages with modified genomes. Nat Microbiol 11, 2336–2348 (2026). 10.1038/s41564-026-02407-2

16. Ceyssens, P.J., Mesyanzhinov, V., Sykilinda, N., et al. The genome and structural proteome of YuA, a new Pseudomonas aeruginosa phage resembling M6. J Bacteriol 190, 1429–1435 (2008). 10.1128/JB.01441-07

17. Lee, Y.J., Dai, N., Walsh, S.E., et al. Identification and biosynthesis of thymidine hypermodifications in the genomic DNA of widespread bacterial viruses. Proc Natl Acad Sci U S A 115, E3116–E3125 (2018). 10.1073/pnas.1714812115

18. Belogurov, A.A., Delver, E.P., Rodzevich, O.V. Plasmid pKM101 encodes two nonhomologous antirestriction proteins (ArdA and ArdB) whose expression is controlled by homologous regulatory sequences. J Bacteriol 175, 4843–4850 (1993). 10.1128/jb.175.15.4843-4850.1993

19. Brinkman, E.K., Chen, T., Amendola, M., van Steensel, B. Easy quantitative assessment of genome editing by sequence trace decomposition. Nucleic Acids Res 42, e168 (2014). 10.1093/nar/gku936

20. Su, T., Liu, F., Chang, Y., et al. The phage T4 DNA ligase mediates bacterial chromosome DSBs repair as single component non-homologous end joining. Synth Syst Biotechnol 4, 107–112 (2019). 10.1016/j.synbio.2019.04.001

21. Lehman, I.R. DNA ligase: structure, mechanism, and function. Science 186, 790–797 (1974). 10.1126/science.186.4166.790

22. Tomkinson, A.E., Vijayakumar, S., Pascal, J.M., Ellenberger, T. DNA ligases: structure, reaction mechanism, and function. Chem Rev 106, 687–699 (2006). 10.1021/cr040498d

23. Williamson, A., Leiros, H.K.S. Structural insight into DNA joining: from conserved mechanisms to diverse scaffolds. Nucleic Acids Res 48, 8225–8242 (2020). 10.1093/nar/gkaa307

24. Chai, R., Zhang, Q., Wu, J., et al. Single-stranded DNA-binding proteins mediate DSB repair and effectively improve CRISPR/Cas9 genome editing in Escherichia coli and Pseudomonas. Microorganisms 11, 850 (2023). 10.3390/microorganisms11040850

25. Lavigne, R., Burkal’tseva, M.V., Robben, J., et al. The genome of bacteriophage phiKMV, a T7-like virus infecting Pseudomonas aeruginosa. Virology 312, 49–59 (2003). 10.1016/s0042-6822(03)00123-5

26. Rokyta, D., Badgett, M.R., Molineux, I.J., Bull, J.J. Experimental genomic evolution: extensive compensation for loss of DNA ligase activity in a virus. Mol Biol Evol 19, 230– 238 (2002). 10.1093/oxfordjournals.molbev.a004076

27. Kulakov, L.A., Ksenzenko, V.N., Shlyapnikov, M.G., et al. Genomes of “phiKMV-like viruses” of Pseudomonas aeruginosa contain localized single-strand interruptions. Virology 391, 1–4 (2009). 10.1016/j.virol.2009.06.024

28. Wang, J., Liu, F., Su, T., et al. The phage T4 DNA ligase in vivo improves the survival-coupled bacterial mutagenesis. Microb Cell Fact 18, 107 (2019). 10.1186/s12934-019-1160-7

29. Lavigne, R., Roucourt, B., Hertveldt, K., Volckaert, G. Characterization of the bacteriophage PhiKMV DNA ligase. Protein Pept Lett 12, 645–648 (2005). 10.2174/0929866054696127

30. Johnston, J.V., Nichols, B.P., Donelson, J.E. Distribution of “minor” nicks in bacteriophage T5 DNA. J Virol 22, 510–519 (1977). 10.1128/JVI.22.2.510-519.1977

31. Marino, N.D., et al. Discovery of widespread type I and type V CRISPR-Cas inhibitors. Science 362, 240–242 (2018). 10.1126/science.aau5174

