## Supplementary material for "A ΦKMV ligase-dependent DNA repair mechanism that mitigates DNA-targeting nucleases": Figures

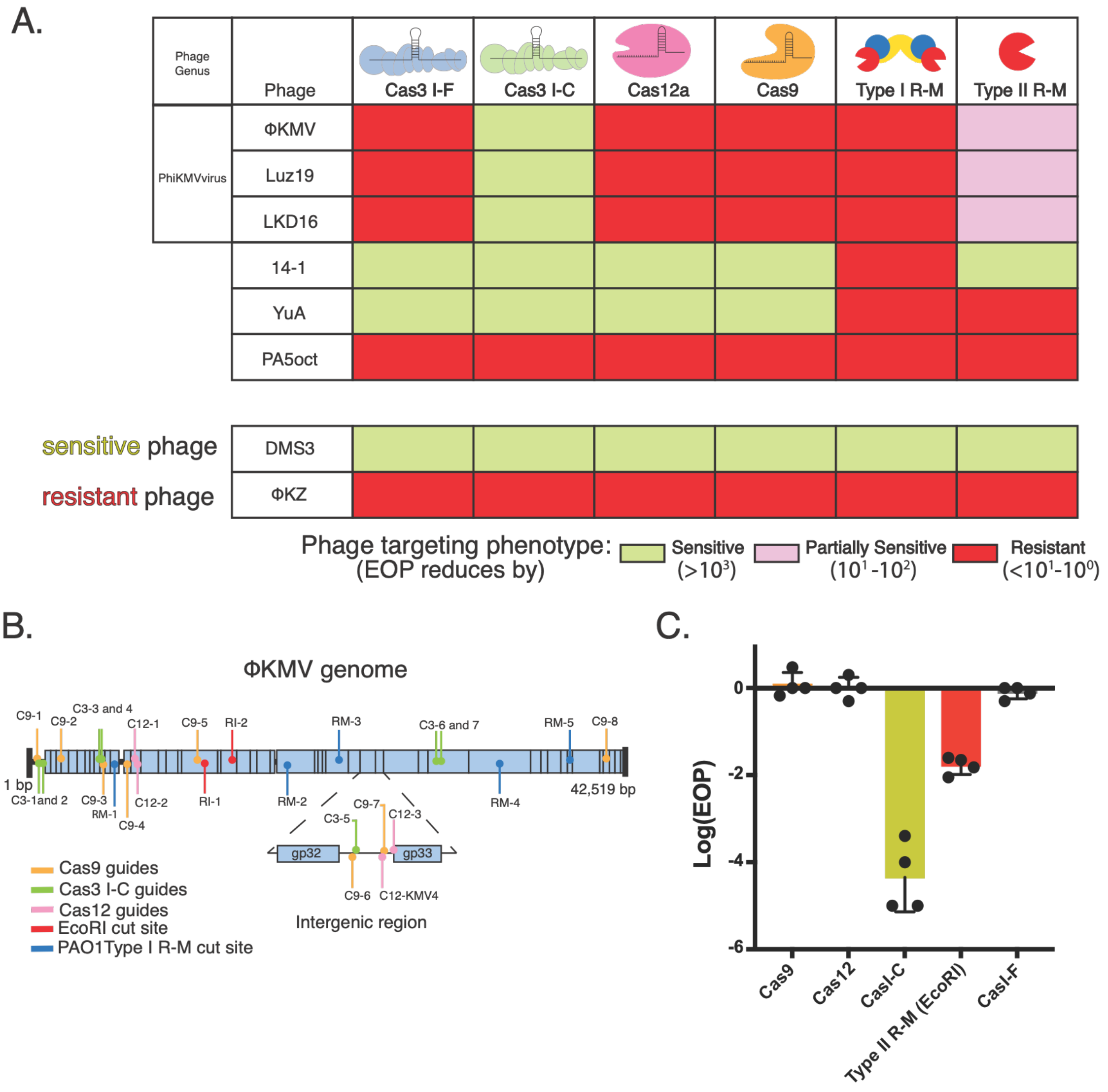


**Figure 1. ΦKMV phages are broadly resistant to native and non-native nucleases**

A. *Pseudomonas aeruginosa* phages were challenged with CRISPR-Cas and Restriction-Modification systems. The phage is denoted as sensitive to the nuclease if efficiency of plating (EOP) is reduced by $>10^3$-fold. B. A schematic depicting all the protospacers used to target ΦKMV with the different DNA targeting systems. C. Log efficiency of plating (EOP) for ΦKMV, as the ratio of phage titer on a targeting strain relative to a non-targeting strain.

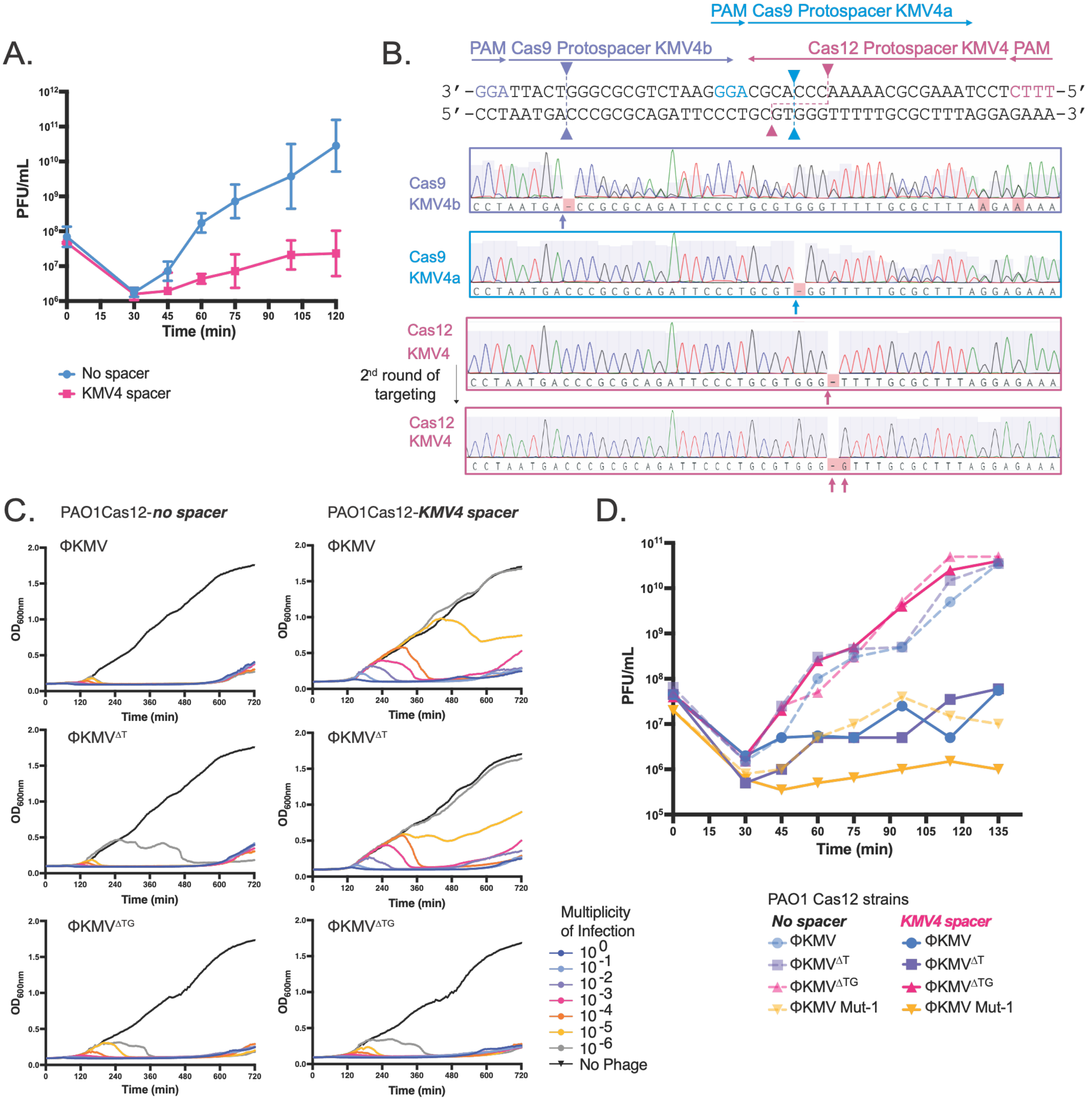


**Figure 2. Small deletions at the cleavage site emerge in the ΦKMV upon Cas12a exposure**

A. Single-step growth curves showing plaque-forming units per mL (PFU/mL) of ΦKMV over time. *Pseudomonas* PAO1 Tn7::Cas12 strains carrying either no spacer (blue) or the KMV4 spacer (pink) were infected with ΦKMV at a MOI of 0.1. Data represent three biological replicates (n = 3). B. ΦKMV intergenic sequence including the Cas9 and Cas12 protospacers. Protospacers for Cas9 KMVb (purple), Cas9 KMVa (blue), and Cas12 KMV4 (pink) are indicated by colored arrows showing spacer orientation and associated PAM sequences. Dashed arrows denote the predicted cleavage sites for each nuclease. Below, corresponding Sanger sequencing traces are shown.

C. Growth curves of PAO1 Cas12 strains infected with ΦKMV, ΦKMV $^{\Delta T}$, or ΦKMV$^{\Delta TG}$ phages across a range of multiplicities of infection (1 to $1 \times 10^{-6}$). Strains expressing either no spacer or KMV4 spacer were infected with phage and optical density at 600 nm ($OD_{600}$) was measured to monitor bacterial growth over time. Data measures technical triplicates. D. Single-step phage growth curves comparing ΦKMV and mutant derivatives, MOI = 1. Dashed lines indicate infections in a no spacer strain, while opaque lines indicate infections in a KMV4 spacer-expressing strain.

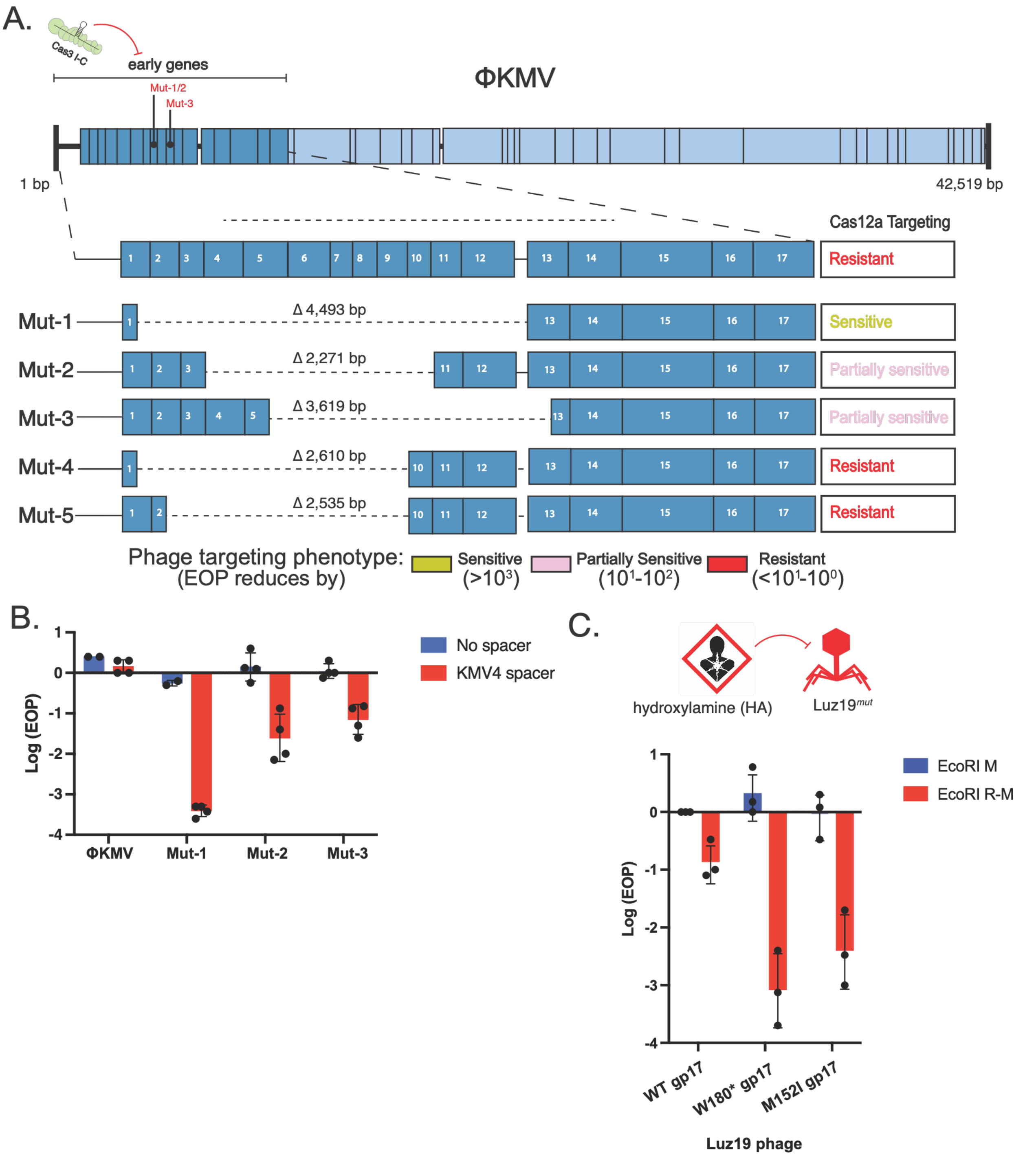


**Figure 3. Mutant phages become sensitized to Cas12a and EcoRI**

A. Schematic of the ΦKMV genome with early genes highlighted in dark blue. Individual early genes were targeted using the type I-C CRISPR-Cas system. ΦKMV mutants 1-3 were generated by targeting *gp8* or *gp10* at the indicated regions. Below, expanded views of each mutant are shown, including the corresponding deletion boundaries. Right column describes phage mutant resistance, sensitivity, or partial sensitivity towards Cas12a. B. Log-transformed efficiency of plating (EOP) for ΦKMV and mutants 1-3 on *PAO1 tn7::cas12* lawns expressing either an empty vector or a KMV4-targeting spacer. C. Schematic of the chemical mutagenesis screen (top). Luz19 phage was treated with hydroxylamine (HA), and mutants were screened for increased sensitivity to the EcoRI restriction-modification (R-M) system. The graph shows log-transformed EOP of wild-type Luz19 and a Luz19 W180* and M152I DNA ligase mutants, on PAO1 strains expressing either EcoRI methylase alone (no restriction; control) or the full EcoRI R-M system. Data represent *n* = 3 biological replicates.

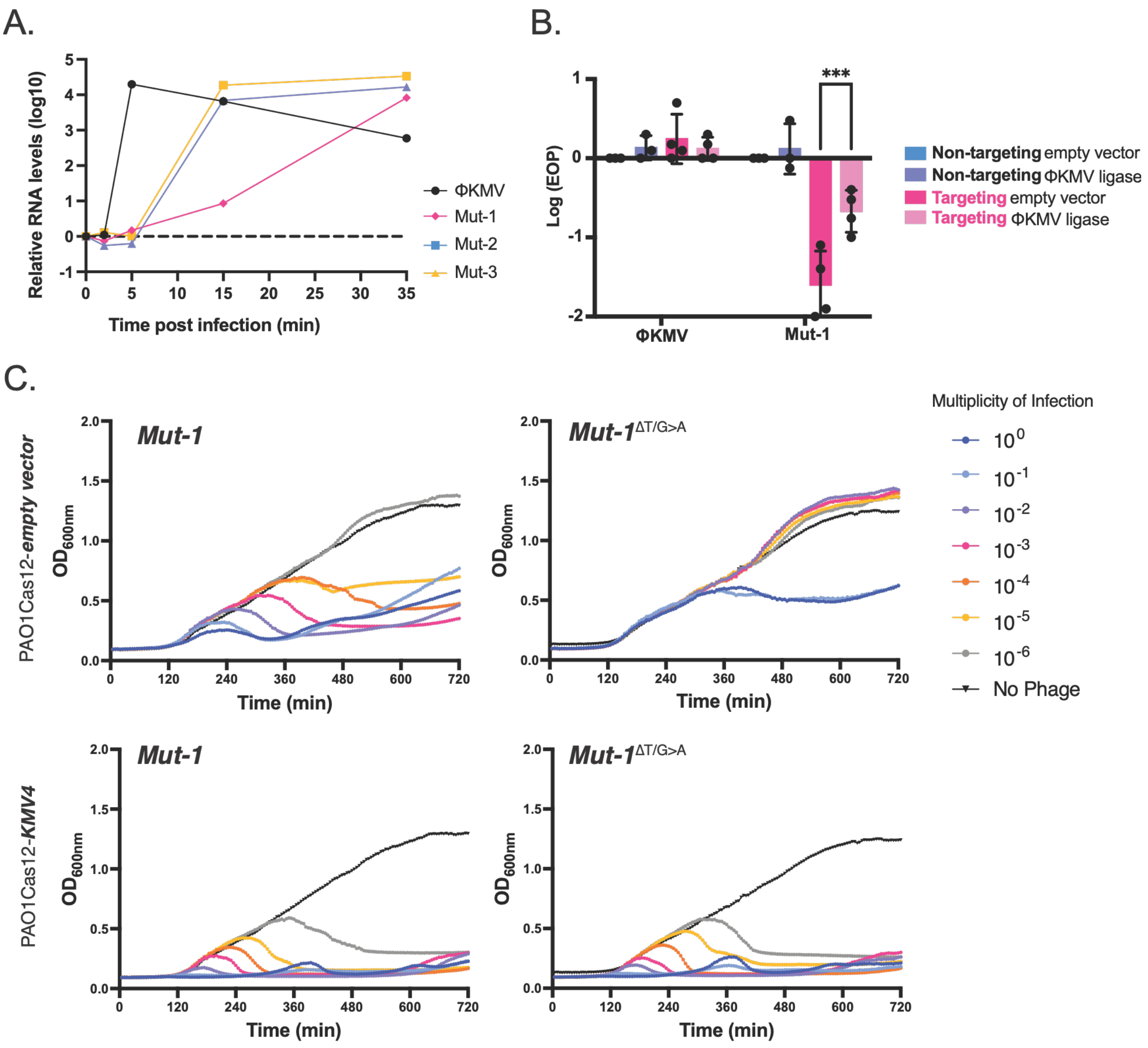


**Figure 4. Delayed ΦKMV ligase expression in Mut-1 is correlated with increased Cas12 targeting**

A. RT-qPCR measurement of gp17 (DNA ligase) over a time course infection. B. *PAO1 tn7::cas12* strains expressing either no spacer or KMV4 spacer, also expressing the ΦKMV ligase. Log EOP is shown for ΦKMV wild-type phage and Mut-1, in biological triplicates (*n=3*). (P-value = 0.0122) C. Growth curves of PAO1 Cas12 strains infected with ΦKMV Mut-1, ΦKMV Mut-1$^{\Delta T/G>A}$ (Mut-1 phage previously targeted at the KMV4 protospacer site) across a range of MOIs (1 to 1×10$^{-6}$). Strains carried either a construct with no spacer or KMV4 spacer. Data represent technical triplicates.

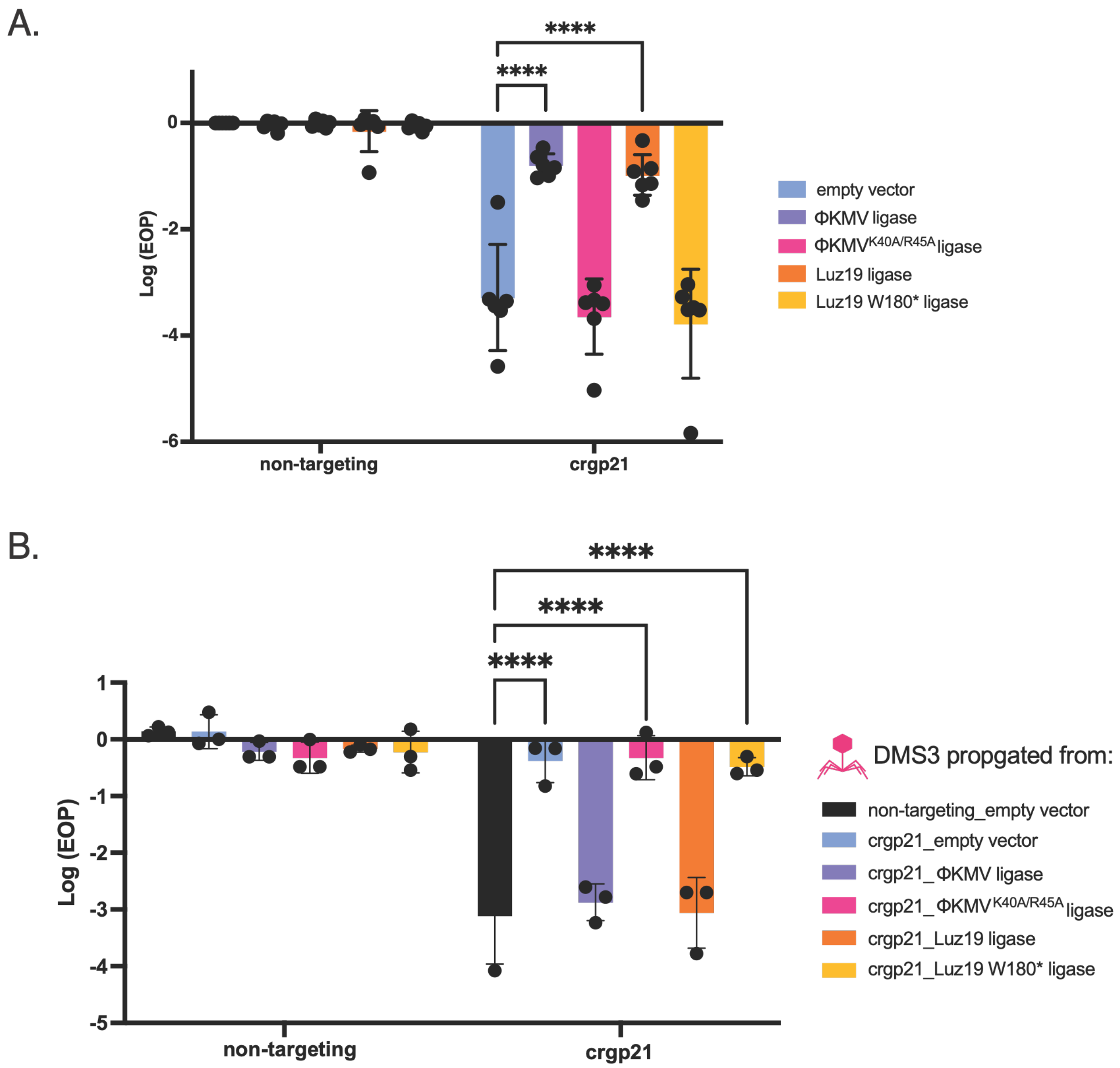


**Figure 5. Expression of ΦKMV and Luz19 ligases rescues Cas12 targeting of DMS3**

A. Full-plate infections of DMS3 phage were performed on *PAO1 tn7::cas12* strains carrying either an empty *attB* site (non-targeting) or *attB::gp21*-targeting spacer. Strains harbored plasmids expressing either no insert (empty vector), ΦKMV / Luz19 ligases, or their corresponding inactive mutants. Log efficiency of plating (EOP) is shown. Data represent $n$ = 5 biological replicates. Statistical significance was determined by two-way ANOVA with multiple comparisons relative to the empty vector control: empty vector vs. ΦKMV ligase, $p < 0.0001$; empty vector vs. Luz19 ligase, $p = 0.0007$. B. DMS3 escaper phages isolated from non-targeting and targeting conditions in the presence or absence of active or inactive ligases. Individual escapers were rechallenged on Cas12-targeting strains. Data represent biological replicates ($n$=2).

**Supplementary Table S1. Strains, phages, plasmids, and crRNAs used in this study.**

| Strains | Plasmid | Description | Reference |
|---|---|---|---|
| PAO1 | N/A | Contains Type I RM system | |
| PAO1 | pSDM160 | pHERD30T with EcoRI Methylase (M) | 10 |
| PAO1 | pSDM161 | pHERD30T with EcoRI-Restriction (RM) | 10 |
| PAO1 | pSG-IF-cas | IPTG inducible, Carb$^R$ pMMBHE plasmid with all Type I-F cas genes (cas1,cas3,csy1-4) | 10 |
| PAO1 | pJB1 | pJW1 with Type II-A sgRNA backbone at the +1 TSS of pBAD | 10 |
| PAO1 *tn7::lbcas12a* | | Chromosomal insertion of Cas12a | 14 |
| PAO1 *tn7::lbcas12a attB::crRNA-DMS3* | | Chromosomal insertion of Cas12a with crRNA targeting DMS3 | 14 |
| LL77 | | Chromosomal insertion of Cas3 Type I-C | 31 |

| Phages |
|---|
| ΦKMV |
| Luz19 |
| LKD16 |
| 14-1 |
| YuA |
| PA5oct |
| DMS3 |
| ΦKZ |

| Plasmids | Description |
|---|---|
| pHERD30T | Empty vector |
| pHERD30T ΦKMV DNA Ligase | For the overexpression of ΦKMV DNA Ligase |
| pHERD30T ΦKMV Ligase K40A/R45A | For the overexpression of mutant ΦKMV Ligase K40A/R45A |
| pHERD30T Luz19 DNA Ligase | For the overexpression of Luz19 DNA Ligase |
| pHERD30T Luz19 Ligase W180* | For the overexpression of mutant Luz19 Ligase W180* |
| pHERD30T crRNA KMV1 | Contains Cas12 spacer targeting ΦKMV /Luz19/LKD16 |
| pHERD30T crRNA KMV2 | Contains Cas12 spacer targeting ΦKMV /Luz19/LKD16 |
| pHERD30T crRNA KMV3 | Contains Cas12 spacer targeting ΦKMV /Luz19/LKD16 |
| pHERD30T crRNA KMV4 | Contains Cas12 spacer targeting ΦKMV /Luz19/LKD16 |
| pCTX2 mini | Empty vector |
| pCTX2 KMV4 | Contains Cas12 target sequence for KMV4 spacer |

| Phage | CRISPR System | Guide Name | Guide Sequence |
|---|---|---|---|
| ΦKMV | Cas3 I-F | I-F G1 | gcaaccatcgacatgtccgaaacctccaccgg |
| ΦKMV | Cas3 I-F | I-F G2 | cgcagattccctgcgtgggtttttgcgcttta |
| ΦKMV | Cas3 I-C | C3-1 | caagcgcggcacaagccataggcgcaagggacac |
| ΦKMV | Cas3 I-C | C3-2 | aatcggcccgccgatactacccagcatactgtca |
| ΦKMV | Cas3 I-C | C3-3 | caccgtggccattccttgccgcgaaccaactcaa |
| ΦKMV | Cas3 I-C | C3-4 | gacggtggccggcttcttgatcagcttggcgccc |
| ΦKMV | Cas3 I-C | C3-5 | cctgcgtgggtttttgcgctttaggagaaaccct |
| ΦKMV | Cas3 I-C | C3-6 | tataccaacatcatgacctccgaggatatcccgc |
| ΦKMV | Cas3 I-C | C3-7 | gtcgccggtggcctgagcctgcacagccttggcc |
| ΦKMV | Cas3 I-C | I-C gp01 | gcaagatgcgtgacgtgcgctacgctaccgaccc |
| ΦKMV | Cas3 I-C | I-C gp02 | cgcaaggctgaacaggcgcaagctaagcagcccc |

| | | | |
|---|---|---|---|
| ΦKMV | Cas3 I-C | I-C gp03_a | cgtggcacagaagagatcaaccgcgctatcgacg |
| ΦKMV | Cas3 I-C | I-C gp03_b | aactgaccggcctgtcgatcatccaccacatcga |
| ΦKMV | Cas3 I-C | I-C gp03_c | aagcctgagaaagcgctggaccaggagtttaacc |
| ΦKMV | Cas3 I-C | I-C gp04 | gtggtagtgggcgccggcgtgaccgtgaaccgct |
| ΦKMV | Cas3 I-C | I-C gp06 | ctcatcgtgaactggagcaagcgtatatgaatcc |
| ΦKMV | Cas3 I-C | I-C gp07 | aggcccgtggcacatccggtccgtggtggttcac |
| ΦKMV | Cas3 I-C | I-C gp08 | caggacgcatgcgtgatcgggtgcatctggccac |
| ΦKMV | Cas3 I-C | I-C gp09 | cggtgagatgtatcgggtagtgcagcccgaggct |
| ΦKMV | Cas3 I-C | I-C gp10 | caccacaccgccctggccaagctgaccgaggtct |
| ΦKMV | Cas3 I-C | I-C gp11 | gcaaccaccaggcgaccatccgcctgttgcaaaa |
| ΦKMV | Cas3 I-C | I-C gp17_a | agtgaggcgaccgggcgtatggcccgcaagaccc |
| ΦKMV | Cas3 I-C | I-C gp17_b | gaggggtcgatggagaaagacccgagcctgacct |
| ΦKMV | Cas12a | KMV1 | tgggaggaatggtaccaatggca |
| ΦKMV | Cas12a | KMV2 | gccaggagtcacggcgcagtgcc |
| ΦKMV | Cas12a | KMV3 | ggagaaaccctatgctactactc |
| ΦKMV | Cas12a | KMV4 | tcctaaagcgcaaaaacccacgc |
| ΦKMV | Cas9 | IIA_G1 | ccaagcgcggcacaagccat |
| ΦKMV | Cas9 | IIA_ G2 | gcttgtgatgagtccagcga |
| ΦKMV | Cas9 | IIA_ G3 | catccgcgccgtccagtcgt |
| ΦKMV | Cas9 | IIA_ G4 | acgccgagaactggtggcag |
| ΦKMV | Cas9 | IIA_KMV-4A | taaagcgcaaaaacccacgc |
| ΦKMV | Cas9 | IIA_KMV-4B | gggaatctgcgcgggtcatt |
| ΦKMV | Cas9 | SG1 | cctccagcagctcccgtcga |
| ΦKMV | Cas9 | SG2 | gcacagcgttcaagcgcttc |

**Supplementary Table 2. Mutant Phages**

A summary of mutant phages generated in this study.

| Phages | Source | Description |
|---|---|---|
| ΦKMV Mut-1 | Generated using Cas 3 Type I-C targeting at the gp08 spacer | Phage sensitive to Cas12; deletion spans gp01 to gp12 |
| ΦKMV Mut-2 | Generated using Cas 3 Type I-C targeting at the gp08 spacer | Phage partially sensitive to Cas12; deletion spans gp04 to gp10 |
| ΦKMV Mut-3 | Generated using Cas 3 Type I-C targeting at the gp10 spacer | Phage partially sensitive to Cas12; deletion spans gp06 to gp13 |
| P2-6 | ΦKMV mutant generated using Cas 3 Type I-C targeting at the gp02 spacer | Phage partially resistant to Cas12; deletion spans gp01 to gp09 |
| P2-7 | ΦKMV mutant generated using Cas 3 Type I-C targeting at the gp02 spacer | Phage partially resistant to Cas12; deletion spans gp02 to gp09 |
| ΦKMV$^{\Delta T}$ | ΦKMV mutant phage after one round of Cas12 targeting at the KMV4 protospacer. | Single T deletion at the predicted Cas12 cut site. |
| ΦKMV$^{\Delta TG}$ | ΦKMV mutant phage after two rounds of Cas12 targeting at the KMV4 protospacer. | T deletion and G insertion at the predicted Cas12 cut site. |
| Luz19 W180* | Generated from chemical mutagenesis | Early stop codon in gp17, DNA ligase; Mutant phage sensitive to EcoRI. |
| Luz19 M152I | Generated from chemical mutagenesis | Mutation in gp17, DNA ligase; Mutant phage sensitive to EcoRI. |

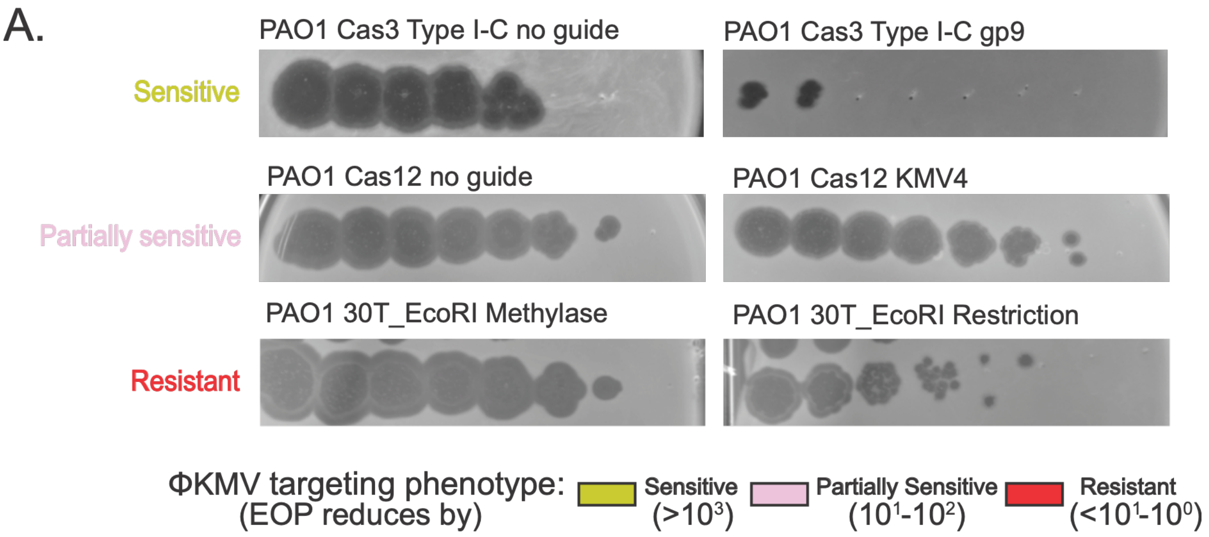


**Supplementary Figure 1 Examples of ΦKMV spot assays used to determine phage sensitivity**

A. Spot Assays measure phage replication of ΦKMV spotted on the lawn of nontargeting and targeting strains. On the left side of the panel, phage phenotype is highlighted as sensitive, partially sensitive, and resistant. Similar spot assays were used to generate the summarized phage phenotypes in Figure 1A.

A.

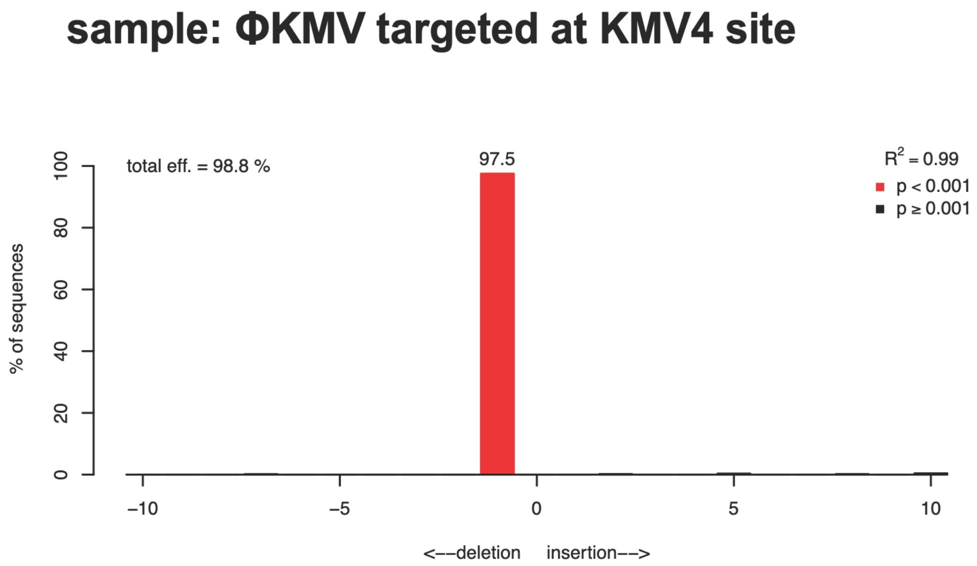


B.

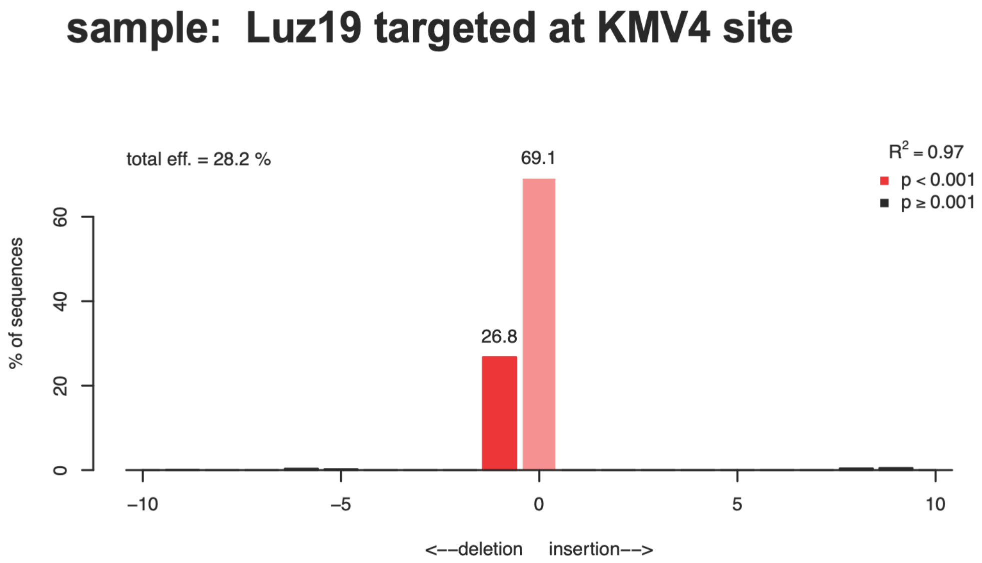


**Supplementary Figure 2 TIDE analysis of a population of ΦKMV and Luz19 phages propagated on PAO1 Cas12 strain with KMV-4 spacer**

A. TIDE analysis of ΦKMV phage targeted at the KMV4 site with CRISPR Cas12. On the y-axis the graph depicts percentage of sequences, total eff or total efficiency is the percentage of sequences that contain indels. The x-axis at 0 represent sequences without indels; +/- n (number on the x axis is the number of indels present in the sequences input). The red bar represents 99.1% of sequences input contain a single deletion at a p value < 0.001. B. TIDE analysis of Luz19 phages.

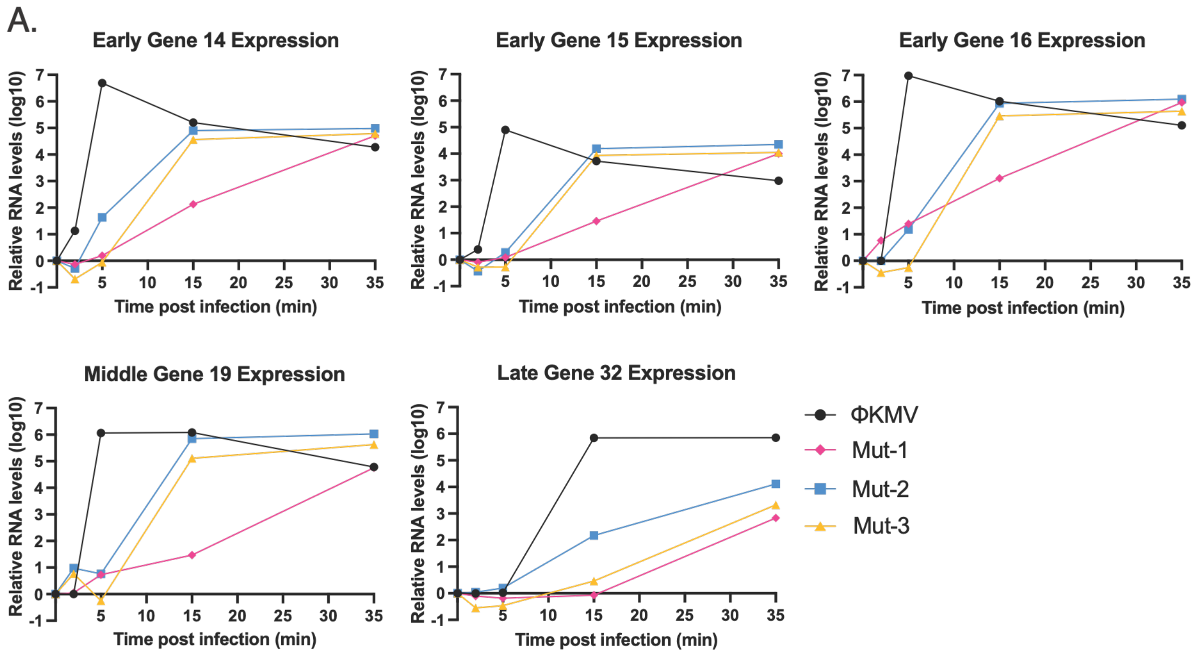


**Supplementary Figure 3 Cas I-C mutants and gene expression profiles of ΦKMV and mutants**

A. Gene expression profiles of middle and late genes measured by RTPCR for ΦKMV (black), and Mut1-3 (yellow, pink, and blue, respectively). The y-axis measures the expression of the target gene against a reference gene. The X-axis measures time post infection with the indicated phages.

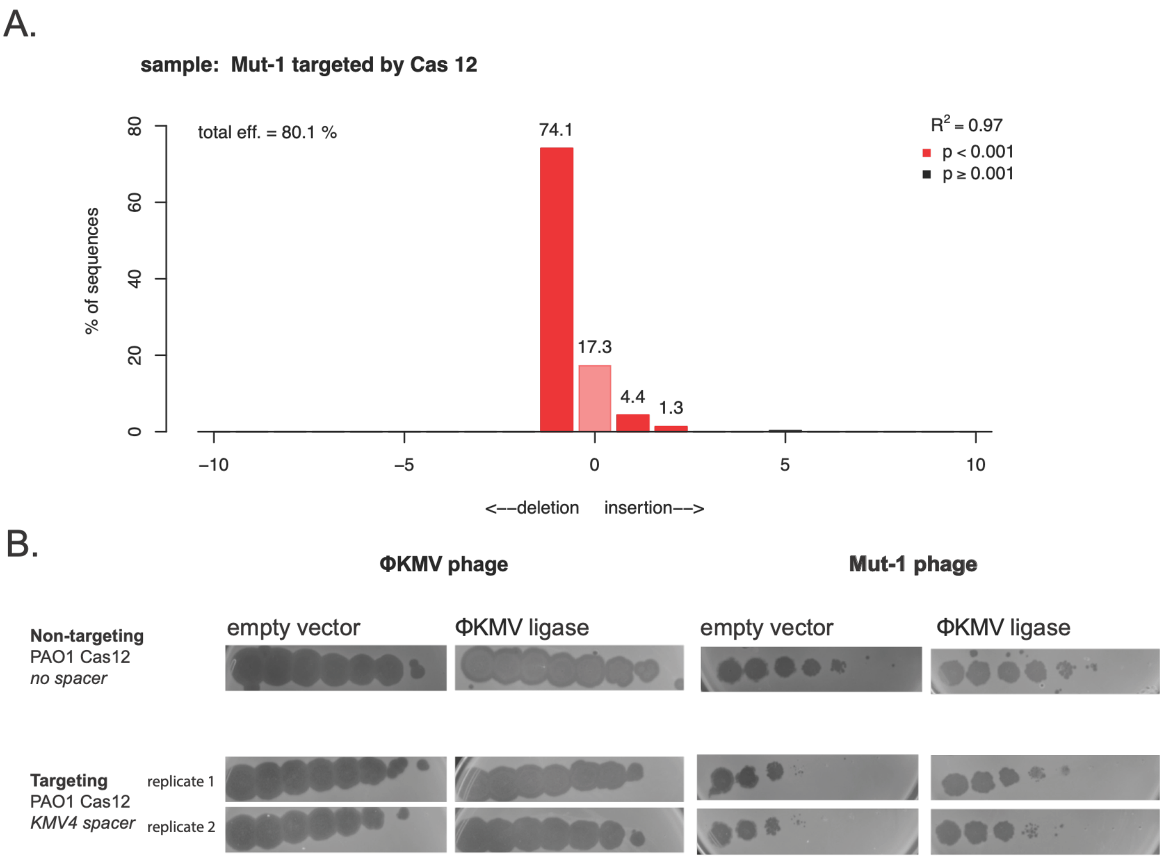


**Supplementary Figure 4 Mut-1 phage which generates many indels after targeting, is complemented by ΦKMV ligase expression**

A. TIDE analysis on ΦKMV Mut-1 phage targeted at the KMV4 protospacer by Cas12. A total efficiency of 80.1% represents the population of sequences with indels. The decomposition of those indels shows 17.3% include no indels, 74. 1% include a single deletion, 4.4% included a single insertion, and 1.3 % included 2 insertions. A wildtype ΦKMV sequence was used as the control reference for TIDE analysis. B. Spot Assays measure phage replication of ΦKMV (on the left) spotted on the lawn of nontargeting and targeting strains, with the extrachromosomal expression of empty vector or active ligase. On the right side of the panel, Mut-1 phage is spotted against the same strains. This data highlights the improved replication of Mut-1 when active ΦKMV ligase is expressed during targeting. The targeting strain spot assays are biological replicates.

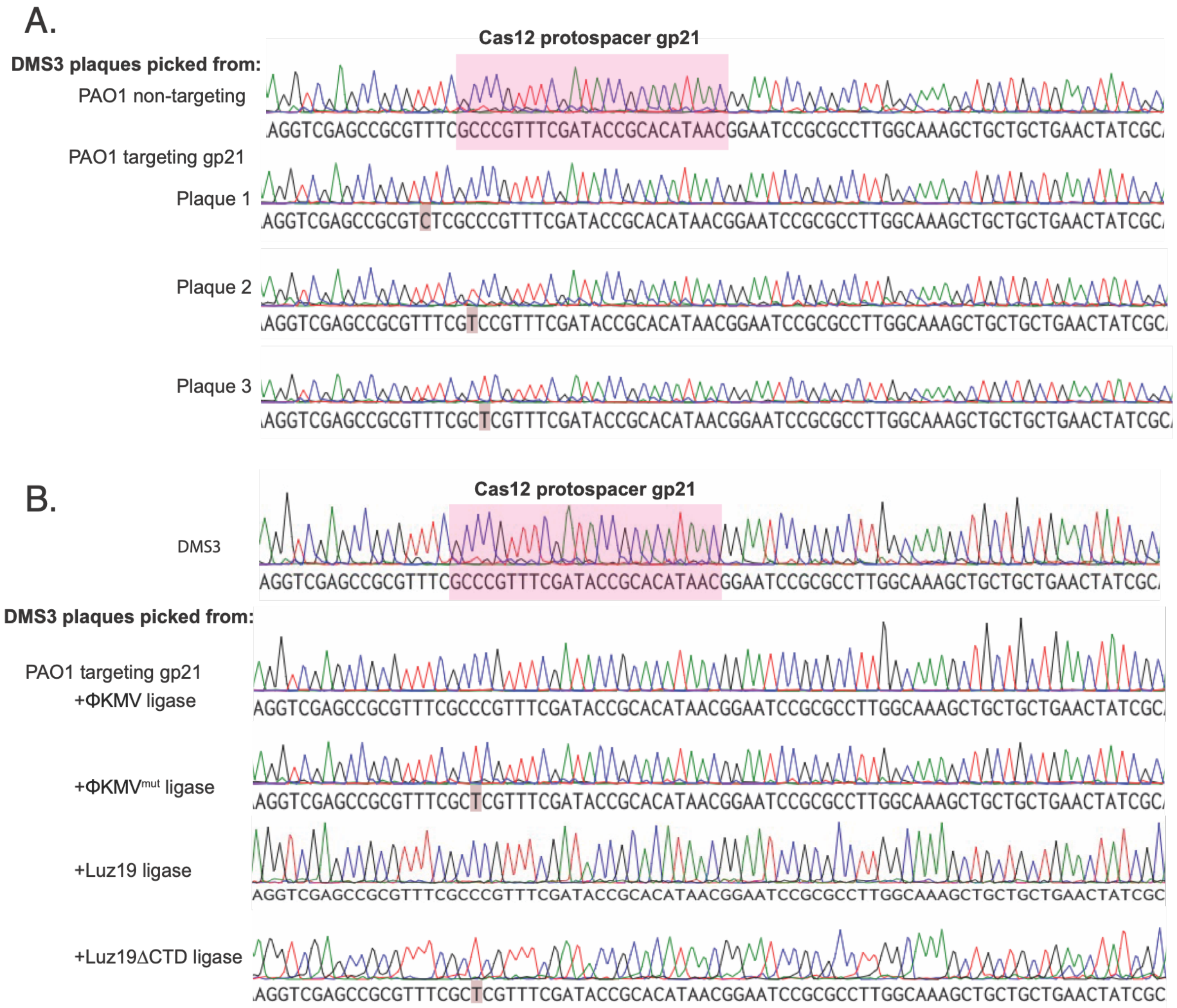


**Supplementary Figure 5 Sanger Sequence traces reveal ΦKMV ligase is sufficient to help DMS3 escape targeting without generation of indels**

A. Sanger Sequencing of region surrounding the gp21 Cas12 protospacer in DMS3. The first read is of a wildtype DMS3 phage picked on a non-targeting PAO1 strain, highlighted in pink is the sequence of the protospacer. The bottom 3 reads are individual plaques picked of DMS3 propagated on a gp21 targeting strain. The sequences here show accumulation of PAM and seed mutations. The PAM sequence which is not highlighted here is the TTTN sequence downstream of the protospacer seen in pink. B. Like Fig S5A, the top sequence is of a wildtype DMS3 phage and the region around the protospacer in pink. The bottom sequences are of DMS3 single plaques picked off a lawn of PAO1 Cas12 targeting (gp21) strains with the added vector of ΦKMV ligase, ΦKMV catalytically dead ligase (K40A/R45A), Luz19 ligase, and Luz19 ΔCTD ligase. Over expression of active ligase produced perfect sequence reads, while the inactive ligases produced mutations in the seed region of the sequence.
